# Wilm’s tumor 1 promotes memory flexibility

**DOI:** 10.1101/101360

**Authors:** Chiara Mariottini, Leonardo Munari, Ellen Gunzel, Joseph M. Seco, Nikos Tzavaras, Jens Hansen, Sarah A. Stern, Virginia Gao, Hossein Aleyasin, Evren U. Azeloglu, Georgia E. Hodes, Scott J. Russo, Vicki Huff, Marc Birtwistle, Robert D. Blitzer, Cristina M. Alberini, Ravi Iyengar

## Abstract

Under physiological conditions, strength and persistence of memory must be regulated in order to produce behavioral flexibility. In fact, impairments in memory flexibility are associated with pathologies such as post-traumatic stress disorder or autism; however the underlying mechanisms that enable memory flexibility are still poorly understood. Here we identified the transcriptional repressor Wilm’s Tumor 1 (WT1) as a critical synaptic plasticity regulator that decreases memory strength, promoting memory flexibility. WT1 was activated in the hippocampus following induction of long-term potentiation (LTP) or learning. WT1 knockdown enhanced CA1 neuronal excitability, LTP and long-term memory whereas its over-expression weakened memory retention. Moreover, forebrain WT1-deficient mice showed deficits in both reversal, sequential learning tasks and contextual fear extinction, exhibiting impaired memory flexibility. We conclude that WT1 limits memory strength or promotes memory weakening, thus enabling memory flexibility, a process that is critical for learning from new experience.

---

Learning produces long-term memory retention and storage by activating molecular mechanisms that consolidate and strengthen an initially labile experience representation. The process of memory strengthening must be regulated in order to remain within the physiological ranges; excessively weak or excessively strong memories are in fact maladaptive and pathological. Weak memories can result from impairments in any of several different processes - storage, retrieval or consolidation (the stabilization process that forms long-term memories) or by an overactive forgetting process ^1-6^ All these processes likely play important roles in memory disorders, in Alzheimer’s disease (AD), aging-related memory loss, and neurodevelopmental cognitive impairments. Conversely, an excessive memory consolidation, and/or impaired forgetting may produce excessively strong and inflexible memories, possibly leading to diseases such as posttraumatic stress disorder (PTSD), autism spectrum disorder (ASD), schizophrenia and obsessive compulsive disorder (OCD). Therefore, the ability to regulate the intensity of memory consolidation and strengthening is of great importance for adaptive behaviors and mental health.

The biological mechanisms required for promoting memory consolidation and strengthening have been investigated in many species and types of memory, identifying roles for a variety of signaling networks ^7-11^, transcription factors ^12-14^and epigenetic changes ^15-17^. However, little is known about mechanisms that reduce memory consolidation and strengthening in order to enable behavioral flexibility. A key question is whether consolidated memories are weakened through a passive decay process, and/or by a learning-induced, active mechanisms that serves to promote memory flexibility. In other words, do signaling pathways that are activated during experience not only support consolidation, but also include counteracting molecular regulators that can decrease memory strength and favor forgetting ^6^, such as the Rho family of GTPases signaling G proteins (Rac) ^18-20^, scribble scaffolds ^21^, DAMB dopamine receptors ^22^, inhibition of AMPA receptor recycling ^22,23^,and neurogenesis ^5,25^?

We therefore tested the hypothesis that memory flexibility results from an active process that occurs in parallel with memory consolidation and strengthening. If this is the case, then mechanisms enabling memory flexibility should be activated and/or induced by learning.

Memory consolidation engages complex regulation of genes transcription activation and repression ^10^. Whereas the role of transcription activators, such as members of the CREB, C/EBP, AP1, NFkB, Rel, Egr 1 and2, and Nurr families have been more extensively documented as promoters of memory consolidation and strengthening ^2,11,13,26-30^, less is known about the role of transcription repressors ^10,31,32^, A few transcription repressors that directly bind to promoter/enhancer DNA sequences in memory formation have been documented: CREB ^33-36^, MeCP2 ^37^, DREAM (downstream regulatory element antagonistic modulator ^38,39^ myocyte enhancer factor-2 (MEF2)^40,41^. The literature thus far suggests that induction of transcription activation correlates with memory strengthening, whereas induction of transcription repression correlates with memory weakening or forgetting ^6,10,31,32^

To search for transcription repressors of plasticity, we screened for transcription repressors activated by induction of LTP at hippocampal excitatory synapses, a cellular model of learning and memory^42^. We identified Wilm’s tumor 1 (WT1), a protein that is important for kidney and gonads development ^43^. Wilm’s tumor 1 is a form of kidney cancer that primarily affects children ages 3 to 4. Interestingly one of the health conditions due to *Wt1* germline mutations is the WAGR syndrome, a disorder characterized by Wilm’s Tumor (W), aniridia (A), genitourinary anomalies (G) and mental retardation (R). Patients with WAGR syndrome have difficulties in learning, processing and responding to information; they may develop behavioral and cognitive abnormalities such as anxiety, obsessive-compulsive disorder (OCD), depression, attention deficit hyperactivity disorder (ADHD) and autism^44^.While the *Wt1* gene has been well characterized for its role in kidney development and function, it has never been investigated in the context of normal brain physiology or cognition. WT1 has been linked to neurodegeneration associated with Alzheimer disease ^45^, and a recent study has showed that during early neuronal development its transcriptional activity is repressed to allow normal neuronal differentiation ^46^. In this study we used different types of genetic and molecular manipulation to investigate the functional role of WT1 in memory consolidation and strengthening and its ability to regulate memory flexibility and new learning. We also investigated WT1 hippocampal physiology by examining the effects of ablating WT1 on pyramidal cell excitability, synaptic plasticity and regulation of entorhinal cortex-hippocampus circuitry. Finally we identified numerous transcriptional targets of WT1 in the hippocampus and functionally characterize one of these genes in plasticity experiments.

## Results

### Learning-induced Wilm’s Tumor 1 (WT1) decreases memory strength

To identify transcription factors activated or induced by long-term plasticity, we employed a protein-DNA binding array assay on rat hippocampal slices in which long-term potentiation (LTP) was induced by strong high-frequency stimulation (Strong-HFS) of the Schaffer Collaterals ^42^. We identified nearly 40 transcription factors whose binding was increased (Fig. 1a and Supplementary Fig. 1a). One of these transcription factors, WT1, is a transcriptional repressor shown to be involved in regulating kidney development ^43^ and in mRNA transport and translation in several cell lines ^47-49^

**Fig. 1:**
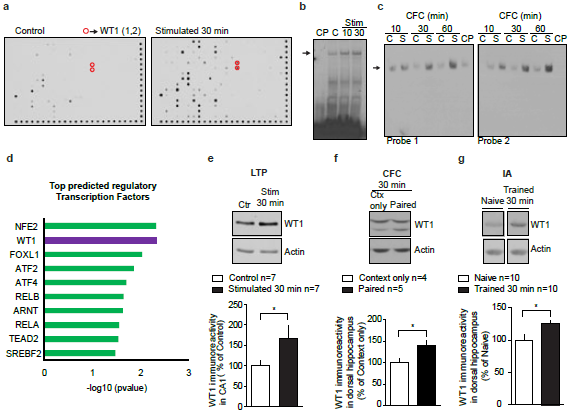
Wilm’s Tumor 1 (WT1) expression and DNA-binding activity are induced by synaptic plasticity and learning. **a**, Protein-DNA binding assay comparing rat hippocampal CA1 extracts from control tissue (left) with extracts obtained from tissue where LTP was induced and collected 30 minutes after stimulation with Strong-HFS (right). WT1 is circled in red; numbers in parentheses indicate two different DNA probes with WT1 consensus sites. **b**, EMSA showed that *in vitro* WT1 binding to a DNA consensus sequence (arrow indicates the WT1/DNA complex) was increased 10 and 30 minutes after stimulation using a protocol that produces L-LTP compared to control unstimulated CA1 extracts (C). The specificity of DNA-protein binding was verified by incubation with excess unlabeled cold probe (CP). **c**, EMSA showed increased WT1 binding to DNA after CFC (arrow indicates the WT1/DNA complex). Nuclear proteins from dorsal hippocampi were extracted from rats where the context was paired to foot shocks (3 shocks at 0.7 mA, indicated as “S”) at indicated times after training and compared to nuclear extracts obtained from rats that were exposed to the same context but that did not receive any foot shock (indicated as “C”). The specificity of DNA-protein binding was verified by incubation with excess of unlabeled cold probe (CP). **d**, Bar graph of the top 10 TFs predicted to regulate gene expression profiles in rat tissue obtained 90 minutes after a stimulation that produced LTP. **e**, Expression of WT1 was significantly increased in rat CA1 stratum radiatum 30 minutes after stimulation with an LTP-inducing stimulus (data are expressed as mean ± s.e.m.; paired t test: *p=0.0495; t=2.455, df=6). **f**, 30 minutes after training expression of WT1 was significantly increased in the dorsal hippocampus of rats trained in CFC (3 shocks at 0.7 mA, Paired) compared to rats exposed to the conditioning chamber but not shocked (context only) (data are expressed as mean ± s.e.m.; unpaired t test: *p=0.0385; t=2.543, df=7). **g**, Expression of WT1 was significantly increased in the dorsal hippocampus of rats trained in an IA task. Protein expression was measured 30 minutes after training and compared to naïve animals (data are expressed as mean ± s.e.m.; unpaired t test: *p=0.0187; t=2.583, df= 18).

Strong-HFS as well as contextual fear conditioning (CFC) learning increased the binding of WT1 to its DNA consensus sequence in the hippocampus of rats (Fig. 1b,c), providing functional evidence for an active involvement of WT1 in these functions. Furthermore, we found independent evidence for WT1 activation in mRNA-seq experiments that identified increased expression of transcripts 90 minutes after LTP induction. Enrichment analysis of this transcriptomic data (see Supplementary Table S1 for complete list of differentially expressed transcripts) predicted WT1 as the second most likely candidate to regulate LTP-induced gene expression followed by members of the CREB family (ATF2 and ATF4) (Fig. 1d; see Supplementary Table S2 for predicted transcription factors analysis).

Additionally both LTP induction (LTP, Fig. 1e) as well as contextual fear learning in two independent tasks, contextual fear conditioning (CFC, Fig. 1f) and inhibitory avoidance (IA, Fig. 1g), resulted in significant increases in the expression levels of WT1 protein within 30 minutes.

To determine the functional role of WT1 in memory formation, we knockdown WT1 protein expression using bilateral injections of antisense oligodeoxynucleotides (WT1-AS) into rat dorsal hippocampus and tested the effect on memory retention using two different hippocampal tasks, one aversive (CFC) and one non-aversive (novel object location, NOL; Fig. 2a). As shown, WT1-AS compared to control scrambled oligodeoxynucleotides (SC-ODN) significantly decreased WT1 protein levels in dorsal hippocampus and resulted in a significantly enhanced CFC memory retention 24 hours after training (Fig.2a). Rats injected with either WT1-AS or SC-ODN did not differ in locomotor activity suggesting that the significant difference in CFC freezing was not due to mobility alteration (Supplementary Fig. 2a). Similar results were obtained with NOL, as rats injected with WT1-AS exhibited increased memory at 24 hours after training (Fig. 2a). Furthermore WT1-AS injected rats showed short-term memory retention, at 1 hour after training, comparable to SC-ODN-injected controls (Fig.2a), indicating that WT1 in the hippocampus selectively affects long-term memory. The WT1-AS or SC-ODN groups did not exhibit any difference in total object exploration time (Supplementary Fig. 2b). These findings, based on two distinct hippocampus-dependent tasks, suggest that WT1, whose expression and DNA binding activity increase following training, decreases memory retention.

**Fig. 2:**
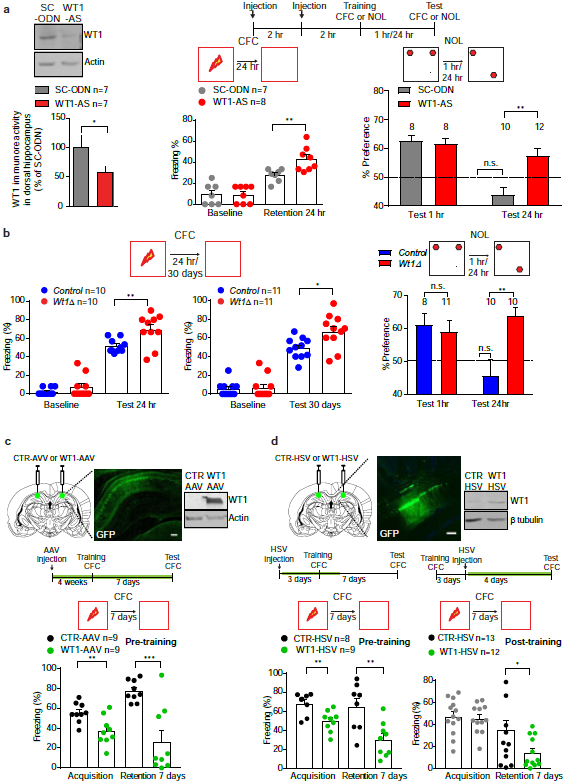
WT1 represses long-term memory consolidation. **a,** Representative western blot and bar graph for the change in WT1 expression obtained with a double injection protocol (2 nmoles/each, 2 hr apart) of WT1-AS (left panel: data are expressed as mean ± s.e.m.; paired t test: *p=0.0362; t=2.687, df=6). Schematic representation of behavioral experiments using WT1 acute knockdown in rats (top panel). Arrows indicate bilateral injections (2 nmol/side) of either WT1-AS or SC-ODN. Injections of WT1-AS increased freezing time in rats trained in CFC and tested 24 h after training (center panel: data are expressed as mean ± s.e.m.; unpaired t test **p=0.0086; t=3.093, df= 13). Injections of WT1-AS did not affect memory retention in NOL 1 hr after training, since both groups showed similar exploration times of the new location (right panel; data are expressed as mean ± s.e.m.; unpaired t test p=0.1685; t=1.438, df= 17. Dashed line indicates 50% preference. Numbers above columns indicate number of animals for each group). In contrast, 24 hr after training WT1-AS-injected animals have better memory than SC-ODN injected ones (right panel: data are expressed as mean ± s.e.m.; unpaired t test: ** p=0.0011; t=3.812, df=20). **b**, *Wt1Δ* animals showed enhanced freezing 24 hr and 30 days (left panel) after training in CFC (data are expressed as mean ± s.e.m.; unpaired t test: for 24 hr, **p=0.0088; t=2.937, df= 18; for 30 days, *p=0.0104; t=2.830, df=20). Both *Control* and *Wt1Δ* mice showed a significant preference for the new location when tested 1 hr after training while only the *Wt1Δ* group showed significant preference for the new location when tested 24 hr after training (right panel; data are expressed as mean ± s.e.m.; unpaired t test: **p=0.0033; t=3.391, df= 18. Dashed line indicates 50% preference. Numbers above columns indicate number of animals for each group). **c**, Immunostaining showing WT1 over-expression via AAV virus 4 weeks after bilateral injection in rats. Scale bar=200 μm. Representative immunoblot showing increase in WT1 expression in the dorsal hippocampus after AAV injection (top panel). Schematic representation of the behavioral experiments using exogenous WT1 expression via WT1-AAV virus in rats. Green highlight line indicates time window for AAV infection and consequent expression of exogenous WT1. Rats were injected with either WT1-AAV or CTR-AAV approximately 4 weeks before training in CFC to allow for maximal over-expression of exogenous protein. Rats injected with WT1-AAV showed significantly reduced levels of freezing during acquisition (comparison between levels of freezing after 3rd shock; data are expressed as mean ± s.e.m.; unpaired t test **p=0.0061; t=3.156, df= 16) and when tested 7 days after training (data are expressed as mean ± s.e.m.; unpaired t test ***p=0.0004; t=4.401, df= 16) compared to CTR-AAV injected rats. **d**, Immunostaining showing WT1 over-expression via HSV virus 3 days after bilateral injection in rats. Scale bar=200 μm. Representative immunoblot showing increase in WT1 expression in the dorsal hippocampus after HSV injection (top panel). Schematic representation of 2 different behavioral experiments using exogenous WT1 expression via WT1-HSV virus in rats. Green highlight line indicates time window for HSV infection and consequent expression of exogenous WT1. HSV injection pre-training: rats injected with WT1-HSV before training, showed a significant difference in their freezing compared to CTR-HSV injected rats both during acquisition (comparison between levels of freezing after 3rd shock; data are expressed as mean ± s.e.m.; unpaired t test **p=0.0042; t=3.377, df= 15) and when tested 7 days after training (data are expressed as mean ± s.e.m.; unpaired t test: **p=0.0049; t=3.299, df= 15). HSV injection post-training: rats were trained in CFC, randomized and injected 3 days after training with either CTR-HSV or WT1-HSV. They were tested 4 days after the injection (total 7 days after training). Rats injected with WT1-HSV showed significantly reduced levels of freezing compared to CTR-HSV injected rats when tested 7 days after training in CFC (unpaired t test: *p=0.0444; t=2.126, df=23).

To extend the investigation of the role of WT1 on synaptic plasticity and memory to different species, we generated genetically modified mice with forebrain expression of an inframe internal *Wt1* deletion, which produces a truncated WT1 protein that lacks zinc fingers 2 and 3 (*Wt1^*fl*/*fl*^; Camk2a-Cre* mice, referred thereafter as *Wt1Δ* mice, Supplementary Fig. 3a). These protein domains are essential for WT1 DNA and RNA binding activity^50,51^ (see methods).

**Fig. 3:**
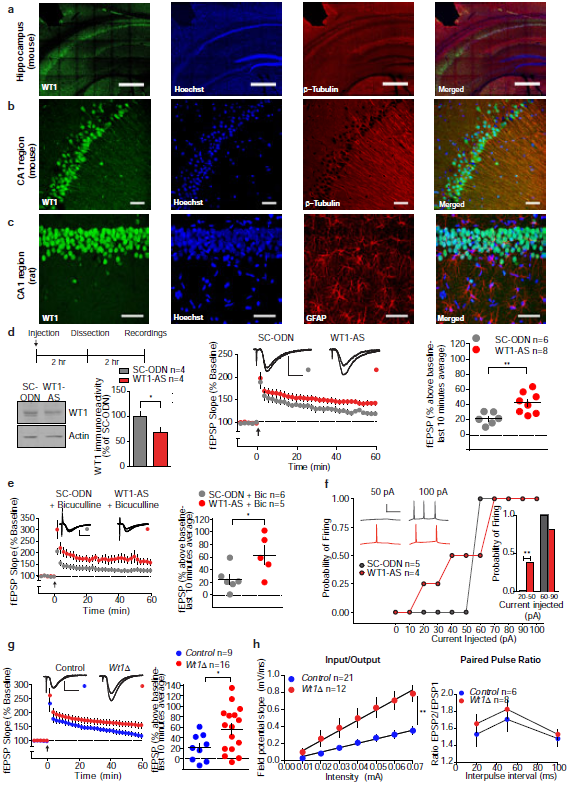
WTl effect is mediated by enhanced activity and excitability of CA1 pyramidal neurons. **a,** Immunostaining of the mouse whole hippocampus shows WT1 localization predominantly within the cell bodies layer. Scale bar=500 μm. **b,** Immunostaining of the mouse CA1 region shows WT1 expression mainly in cell bodies but also in proximal dendrites stained with β-tubulin antibody. Scale bar=50 μm. **c,** Immunostaining of the rat CA1 region shows that WT1 is expressed in pyramidal neurons and not GFAP (glial fibrillary acidic protein) positive astrocytes. Scale bar=50 μm. **d,** Summary scheme for the electrophysiology experiments and representative western blot and bar graph (left panel) for the change in WT1 expression obtained with a single intrahippocampal injection (2 nmoles) of WT1-AS (data are expressed as mean ± s.e.m.; paired t test: *p=0.0202; t=4.521, df=3). Antisense-mediated knockdown of WT1 increased the ability of weak stimulus (delivered at arrow) to induce LTP. Representative fEPSPs show superimposed traces recorded during baseline and 60 min post-HFS. Calibrations: 0.5 mV/10 ms (center and right panel). The increase in fEPSP slope above baseline during the final 10 minutes was greater in slices from hippocampi of rats injected with WT1-AS compared with controls injected with scrambled ODN (data are expressed as mean fEPSP ± s.e.m.; two-way ANOVA RM: F_(1,12)_=10.58, **p=0.0069). **e,** WT1-AS mediated LTP enhancement was not due to effects on inhibitory interneurons, since it was intact in the presence of bicuculline (10 μM). Representative fEPSPs show superimposed traces recorded during baseline and 60 min post-HFS (delivered at arrow). Calibrations: 0.5 mV/10 ms. fEPSP slope over the final 10 minutes of recording showed that treatment with WT1 ODNs significantly enhanced L-LTP (data are expressed as mean fEPSP ± s.e.m.; two-way ANOVA RM: F(1,9)=6.039, *p=0.0363). **f,** WT1 depletion increased excitability of CA1 pyramidal neurons in rat. Whole-cell patch recordings were obtained in current clamp mode, and the number of spikes evoked by a series of depolarizing current steps was counted. Inset on the right shows the probability of evoking at least one spike in response to a weak (20-50 pA) or a stronger (60-90 pA) current step in neurons from WT1-depleted (red) or control (gray) hippocampi (n=4 for WT1-AS, n=5 for SC-ODN; two-tailed Chi-square test, **p=0.0041). Resting membrane potential and input resistance measured -63.75 ±3.15 mV and 105.8 ± 21.76 MΩ in the WT1-AS group, and -60.80 ± 2.85 mV and 109.3 ± 20.04 MΩ in the SC-ODN group. Inset on the left shows representative traces in cells from WT1-AS (red) or SC-ODN (gray) hippocampi. Calibration: 50 mV/100 ms. **g,** Hippocampal slices from *Wt1Δ* mice upon weak stimulus (delivered at arrow) showed enhanced LTP compared to control littermates (*Control*). Representative fEPSPs show superimposed traces recorded during baseline and 60 min post-HFS. Calibrations: 0.5 mV/10 ms. Summary of the final 10 minutes of recording showed that LTP was significantly enhanced in *Wt1Δ* mice compared to their control littermates (data are expressed as mean fEPSP ± s.e.m; two-way ANOVA RM: F(i 23)=5.125, *p=0.0333). **h,** *Wt1Δ* mice showed increased basal synaptic efficiency (left panel: input/output; linear regression unpaired t test, **p=0.0077, t=2.845, df=32) but did not affect paired-pulse ratio (right panel; paired pulse ratio; two-way ANOVA RM, p=0.0878).

*Wt1Δ* mice were viable, of normal size and weight, and did not show any gross alteration in hippocampal morphology compared to wild-type littermates (referred thereafter as *control* mice; Supplementary Fig. 3b). The transgenic mice also were similar to control mice with respect to protein levels in peripheral tissue, as well as in their urine and blood chemistry (metabolic enzyme and electrolyte panel; Supplementary Fig. 3c and Supplementary Fig. 3d).

Similar to rats in which WT1 was knocked down in the hippocampus, *Wt1Δ* mice compared to control mice showed enhanced memory retention 24 hours as well as 30 days after CFC training (Fig. 2b). They also showed enhancement in NOL retention 24 hours, but not 1 hour following training (Fig.2b). The open field activity, pain response and total object exploration time of *Wt1Δ* mice were similar to those of control mice (Supplementary Fig. 4a-c), indicating that the effect of the genotype on NOL and CFC were not due to changes in locomotor activity, pain sensitivity or exploration. In contrast, when tested in the elevated plus maze, a paradigm used to measure anxiety-like behavior, *Wt1Δ* mice spent significantly more time in the closed arm and made a significant lower numberof entries in the open arm (Supplementary Fig. 4d), compared to controls, suggesting that forebrain deletion of WT1 may affect also anxiety behavior regulation.

**Fig. 4:**
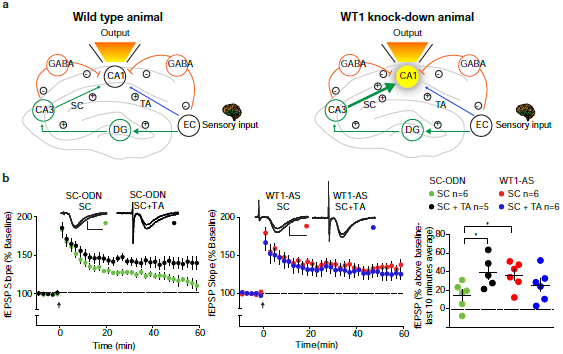
Circuit mechanism of WT1 action. **a,** Schematic representation of WT1 depletion effect on corticohippocampal input to CA1. Left panel (wild type animal): normally, activation of both the direct temporoammonic pathway (blue) and the trisynaptic pathway (green) are required for LTP induction at the Schaffer collateral (SC)**→** CA1 synapse. Right panel (WT1 knock-down animal): in WT1-depleted hippocampus, enhanced basal efficiency of SC**→**CA1 signaling and / or CA1 excitability enable trisynaptic pathway activity alone to induce LTP. EC= entorhinal cortex; DG = dentate gyrus; TA=temporoammonic pathway. **b**, Theta burst stimulation (TBS, delivered at arrow) of the schaffer collaterals (SC) induced stable LTP in slices from rats injected with SC-ODN onlywhen combined with phase-delayed TBS at the temporoammonic (TA) pathway (left and right panels). Conversely, in slices fromWT1-AS-injected hippocampi, the same TBS of SC alone induced LTP, which did not differ from that induced by dual-pathway TBS (center and right panels). Representative fEPSPs show superimposed traces recorded during baseline and 60 min post-TBS. Calibrations: 0.5 mV/10 ms. Data are expressed as mean fEPSP ± s.e.m. Statistical analysis by two-way ANOVA RM: Schaffer collateral stimulation (SC) comparing SC-ODN vs WT1-AS ODNs: F(_1,10_)=6.931, *p= 0.0250; SC-ODN comparing Schaffer collateral stimulation (SC) vs Schaffer collateral stimulation + temporoammonic stimulation: (SC+TA) F(_1,9_)=7.112, *p=0.0258. No significant effect was observed when comparing WT1-AS SC vs WT1-AS SC+TA: F(_1_,_10_)=1.437, p=0.2582.

Collectively these results indicate that the expression and functional activation of the transcriptional repressor WT1 is increased in the hippocampus by learning and that it acts to suppress memory.

To test WT1 function as memory suppressor, we over-expressed wild type WT1 using either an AAV or HSV virus injected into the dorsal hippocampi of rats (WT1-AAV or WT1-HSV respectively; Fig. 2c and 2d). AAV-GFP or HSV-GFP viruses were used as controls (CTR-AAV or CTR-HSV). Rats bilaterally injected into the hippocampus with either viruses were trained either 4 weeks (AAV) or 3 days (HSV) after infection, times that correspond to the respective peaks of viral expression for the two viruses ^52^. As shown in Fig. 2c and Fig. 2d (pre-training) both WT1-AAV-and WT1-HSV-injected rats had a significantly decreased memory retention 7 days after CFC training. These data indicate that WT1 over-expression is sufficient for reducing memory retention. Notably, because over-expression of WT1 significantly reduced the acquisition of the task (Fig. 2c and Fig. 2d), we tested the effect of viral injection following training (post-training). As shown in Fig. 2d (post-training), WT1-HSV compared to control virus decreased memory retention tested 7 days after training. Together these data indicate that over-expression of WT1 is sufficient for decreasing memory retention.

### WT1 modulates plasticity by controlling the excitability of hippocampal CA1 neurons

Immunohistochemical staining of rat and mouse hippocampi obtained from naïve animals revealed that WT1 is predominantly localized within the nuclei of pyramidal neurons (Fig. 3a-c) with a weaker immunoreactivity in the proximal apical dendrites (Fig. 3b). WT1 immunoreactivity was not detected in astrocytes marked by glial fibrillary associated protein (GFAP-positive) (Fig. 3 c).

Given that the effect of decreasing WT1 expression in the hippocampus enhances memory retention, here we tested whether WT1 knockdown affects hippocampal LTP induction and/or maintenance. Western blot analyses showed that single intrahippocampal injection of WT1-AS significantly decreased WT1 levels in hippocampal slices compared to control slices injected with SC-ODN (Fig. 3d). This WT1 knockdown did not alter basal synaptic transmission (Supplementary Fig. 5a), nor did it affect the induction or maintenance of LTP elicited by Strong-HFS (Supplementary Fig. 5b). However, a role for WT1 in synaptic plasticity emerged at synapses activated with a weak high-frequency stimulation (Weak-HFS) protocol, which produced decremental potentiation in control slices but stable LTP in slices from animals injected with WT1-AS (Fig. 3d).

**Fig. 5:**
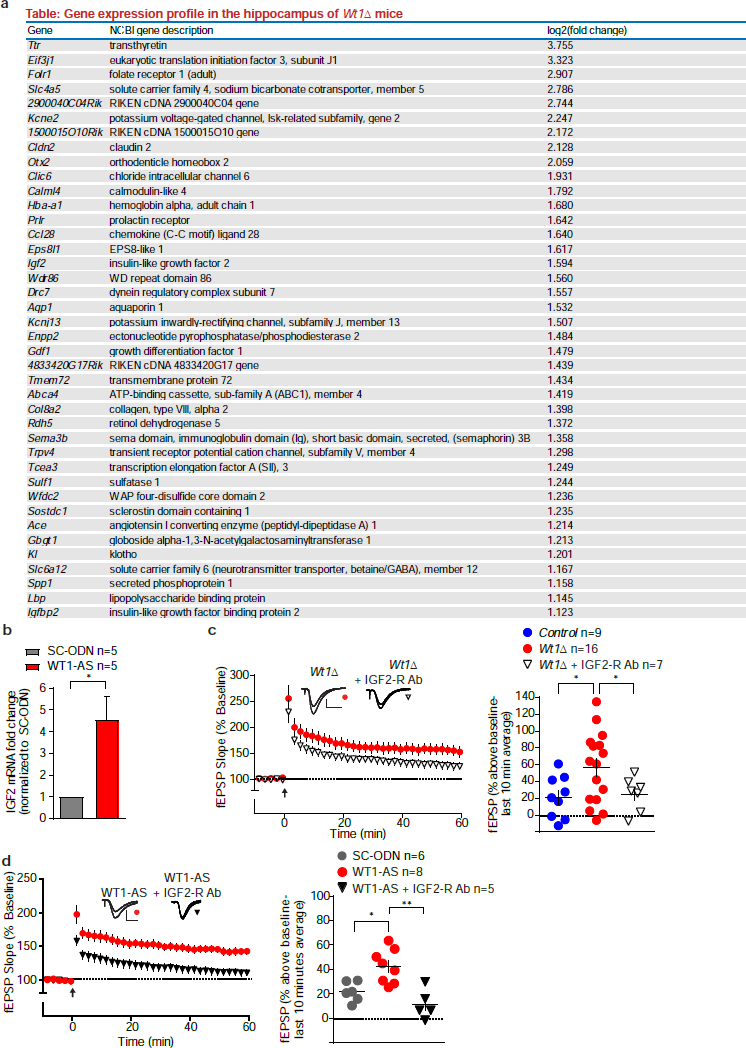
WT1 effects on hippocampal plasticity are mediated via IGF2. **a,** Table representing the top 40 differentially expressed genes whose mRNA expression was regulated in the *Wt1Δ* mice. Numbers indicate the log_2_-fold change for each gene comparing *Wt1Δ* hippocampi with hippocampi from their *Control* wild type littermates. For a list of all the differentially expressed genes, their gene symbols as well as their extended names see Supplementary Table S3. **b**, Quantitative real time PCR showed that WT1 acute knockdown in rats significantly increases IGF2 mRNA expression (data are expressed as mean ± s.e.m.; unpaired t test * p=0.0260; t=3.454, df=4). **c**, The enhanced LTP induced by Weak-HFS in *Wt1Δ* slices was abolished by bath application of a specific antibody against the IGF2 receptor (IGF2-R Ab, 5 μg/ml). Representative fEPSPs show superimposed traces recorded during baseline and 60 min post-HFS. Calibrations: 0.5 mV/10 ms. Summary of the final 10 minutes of recording showed that LTP in hippocampal slices from *Wt1Δ* mice was significantly reduced by bath application of IGF2-R Ab (data are expressed as mean fEPSP ± s.e.m; two-way ANOVA RM, * p=0.0430, F(i, 13) =5.03). For ease of comparison data for the *Control* and the *Wt1Δ* group in the bar graph are the same as in Fig. 3g. **d**, WT1-AS-mediated LTP enhancement was blocked by bath application of a specific antibody directed against the IGF2 receptor (IGF2-R Ab, 5 μg/ml). Representative fEPSPs show superimposed traces recorded during baseline and 60 min post-HFS. Calibrations: 0.5 mV/10 ms (left panel). Summary of the final 10 minutes of recording showed that LTP in WT1-AS injected slices was significantly reduced by bath application of IGF2-R Ab o (right panel; data are expressed as mean fEPSP ± s.e.m; two-way ANOVA RM, ** p=0.0017; F(_1_, _11_)=16.85). For ease of comparison data for the scrambled-ODN and the WT1-AS group, in both the time course and bar graph, are the same that have been shown in Fig. 3d.

To assess whether WT1 knockdown might enhance LTP indirectly through an effect on interneuron function ^53^, we stimulated slices with Weak-HFS in the presence of the GABA-A receptor antagonist bicuculline. Under these conditions, slices from WT1 knockdown hippocampi still showed enhanced LTP, indicating that WT1 likely regulates synaptic plasticity through direct effects on pyramidal neurons (Fig. 3e).

We therefore hypothesized that WT1 knockdown might enhance LTP by increasing pyramidal cell excitability, since postsynaptic spiking during stimulation facilitates LTP induction ^54^.To test this hypothesis, whole-cell recordings were obtained from pyramidal neurons in area CA1 of rat hippocampus. In recordings from WT1-AS injected hippocampi, weak depolarizing currents (20-50 pA) were more likely to evoke action potentials than in neurons of scrambled ODN-injected hippocampi (Fig. 3f) indicating that WT1 knockdown increased excitability. In contrast, in response to relatively strong depolarizing currents (70-100 pA), neurons from WT1-AS slices fired significantly *fewer* action potentials than those treated with scrambled ODN (mean number of spikes = 2.1 ± 0.173 and 1.375 ± 0.125, respectively; unpaired t-test: *p=0.0146; t=3.394, df=6. See traces). No significant differences were observed in the amplitude, frequency or inter-event interval in both spontaneous and mEPSCs (Supplementary Fig. 6a and Supplementary Fig.6b).

**Figure 6:**
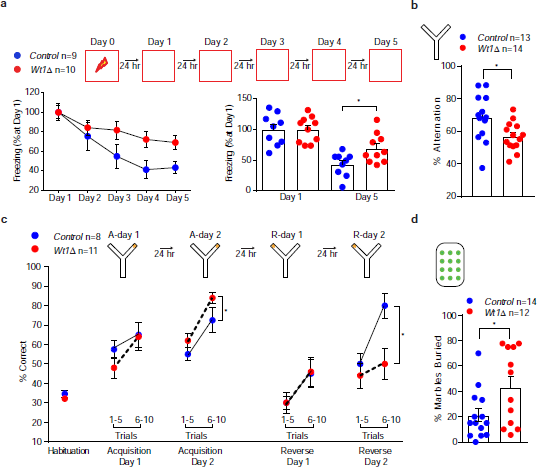
WT1 controls memory flexibility. **a**,*Wt1Δ* mice exhibited a lower rate of extinction than their control littermates measured at day 5 of extinction; % freezing at day 5 is normalized to freezing at day 1 (expressed as mean ± s.e.m.; unpaired t test: *p=0.0196; t=2.578, df=17). **b,** *Wt1Δ* mice showed significant impairment in working memory (% alternation) in the spontaneous alternation task in a Y maze (data are expressed as mean ± s.e.m.; unpaired t test: *p=0.0178; t=2.537, df=25). **c,** *Wt1Δ* mice compared to *Control* littermates showed a significant different performance in the acquisition phase and in the reversal phase when trained in a reversal learning task in a Y-maze. Data are expressed as % correct arm entry (baited arm). A-day 1 and A-day 2: acquisition sessions 1-2; R-day 1 and R-day 2: reversal sessions 1-2 (data are expressed as mean ± s.e.m.; two-way ANOVA RM; *p=0.0348; F_(1,17)_=5.263 for Acquisition phase. *p=0.0120; F_(1,17)_=7.916 for Reversal phase), **d,** *Wt1Δ* mice exhibited a significant difference in repetitive behavior measured via the marble burying test. The bar graph is shows the % of buried marbles during a 30 minutes test (data are expressed as mean ± s.e.m.; unpaired t test: *p=0.0297; t=2.312, df=24).

In agreement with these data in rat hippocampuses, slices from *Wt1Δ* mice also showed sustained hippocampal LTP following Weak-HFS, while slices from control mice produced only transient potentiation (Fig. 3g). When compared to their control littermates, *Wt1Δ* mice showed increased basal Schaffer collateral-CA1 synaptic efficiency with no difference in paired-pulse ratio (Fig. 3h), indicating that WT1 regulates synaptic efficiency through a postsynaptic mechanism.

Collectively, these results suggest that WT1 acts as a synaptic plasticity repressor that dampens the postsynaptic response to a weak stimulus, while preserving the normal dynamic range of the response to superthreshold stimuli.

### WT1 regulates circuitry properties of CA1 pyramidal cells

The role of the CA1 region in memory processing involves the circuit-level integration of information arriving from the entorhinal cortex via two major inputs: (1) the direct temporoammonic (TA) pathway, in which entorhinal neurons of the perforant path synapse on distal apical dendrites of CA1 pyramidal neurons, and (2) an indirect input, in which entorhinal activity provides phase→delayed information to proximal apical dendrites in CA1 through a series of three synapses: perforant path→dentate gyrus, mossy fibers→CA3, and Schaffer collaterals→CA1. The CA1 pyramidal neuron functions as a coincident detector, integrating these temporally segregated streams of information from cortical activity ^55^. This coincidence detection function can be studied in hippocampal slices where the two inputs are activated independently (Fig. 4a; wild type animal) ^56^. We reasoned that WT1 levels could regulate the need for convergent activity of both inputs to induce LTP at the Schaffer collateral (SC)-CA1 synapse. Depletion of WT1 might allow SC stimulation alone to induce LTP without the added information provided by the TA input (Fig. 4a; WT1 knock-down animal). We tested this hypothesis by stimulating CA1 with theta-burst stimulation (TBS) at both the TA and SC inputs, with SC stimulation phase-delayed relative to TA. In hippocampi from control rats injected with scrambled ODN, induction of LTP required activation of both inputs (Fig. 4b). However, the TA input became dispensable in WT1-depleted hippocampus, so that SC stimulation alone was as effective in producing LTP as dual pathway stimulation (Fig. 4b). Thus, the “normal” level of WT1 imposes a requirement for circuit-level computation in the CA1 neuron, leading to LTP. In contrast, in WT1-depleted hippocampus circuit-level computation no longer is necessary: SC→CA1 activity can induce LTP without confirmatory input from TA→CA1. Combined with our findings of increased pyramidal cell excitability and altered spike encoding of depolarization in WT1-depleted hippocampus, this result indicates that WT1 activity regulates the computational properties of CA1 pyramidal cells.

### WT1 target genes: immediate early genes, retinoic acid-related genes, transthyretin, and insulin-like growth factor 2 (IGF-2)

To identify the target genes of WT1 in hippocampal synaptic plasticity and memory, we compared mRNA-seq profiles of *Wt1Δ* and control mice. We identified 193 differentially expressed transcripts (Fig. 5a; see Supplementary Table S3 for complete list).

While transcripts encoding for plasticity and memory related immediate early genes such as the activity-regulated cytoskeletal-associated protein (*Arc*) and the FBJ osteosarcoma oncogene (*Fos*) were significantly downregulated, we found that several genes belonging to the retinoic acid signaling pathway were instead up-regulated. These include retinol dehydrogenase 5 (*Rdh5*), cellular retinoic acid binding protein 2 (*Crabp2*), aldehyde dehydrogenase family 1, subfamily A2 [(*Aldh1a2*; also known as retinaldehyde dehydrogenase 2 (*Raldh2)]* (see supplementary Table S3 for complete list). Interestingly, another upregulated transcript encodes transthyretin (TTR), a protein that is involved with transport of retinol in the plasma and which plays an important role in neuroprotection^57^ as well as memory consolidation and neurogenesis in the hippocampus^58,59^ Furthermore, TTR has also been shown to upregulate hippocampal expression of insulin-like growth factor receptor I (IGF-IR) and its nuclear translocation ^60^.

Notably we found that *Igf2* was ranked sixteenth in our list (and the insulin-like growth factor binding protein 2, known as IGFBP2, was ranked fortieth), as one of the top differentially regulated genes. This is in agreement with previous literature on kidney and cell lines (human fetal kidney and HepG2 cells) reporting that WT1 suppresses the expression of *Igf2*^61,62.^

### IGF-2 mediates WT1 effects on hippocampal synaptic plasticity

In the brain, IGF-2 is required for long-term memory consolidation in the hippocampus, and it has been shown that administration of recombinant IGF-2 significantly enhances memory as well as LTP^63,64.^

Using quantitative real time RT-PCR, we confirmed that acute WT1 knockdown using WT1-AS significantly increased IGF-2 mRNA expression in dorsal hippocampus (Fig. 5b). Based on this finding, we examined whether *Igf2* mediates the enhanced synaptic plasticity produced by WT1-depletion. In hippocampal slices, application of an IGF2 receptor-blocking antibody significantly inhibited LTP enhancement in WT1-deficient mice and rats (Fig. 5 c and 5d) consistent with the hypothesis that, similarly to the kidney, *Igf2* is one of the key downstream targets of the transcriptional repressor WT1. Thus, we conclude that the effects on plasticity observed when WT1 is knocked down or ablated rely on derepression of the *Igf2* gene.

### WT1 enables memory flexibility

A possible role for WT1 is that it limits memory consolidation and strengthening to promote memory flexibility. If this were true, eliminating WT1 function, which results in enhanced CFC, should reduce the ability of the animals to adapt behavioral responses to a changing environment. Thus, we tested whether WT1 depletion affects extinction of CFC memory, reversal learning, repetitive compulsive behavior, and/or sequential learning. Compared to control littermates, *Wt1Δ* mice showed deficient CFC extinction (Fig. 6a), a hippocampal-dependent task by which the animals learns to decrease the conditioned response to fear ^65^. *Wt1Δ* mice also showed reduced spontaneous alternation in a Y maze (Fig. 6b), a paradigm widely used to test active retrograde working memory, based on the general trend of mice to explore the least recently visited arm and thus to alternate their visits among the three arms ^66^. Furthermore, when compared to control littermates, *Wt1Δ* mice showed enhanced memory in the acquisition phase but impairment in the reversal learning phase of the Y maze (Fig. 6c), indicating that WT1 limits the ability to adapt to previously learned responses. Finally, *Wt1Δ* mice also differ from controls in the marble-burying test, a paradigm used to measure repetitive behavior (Fig. 6d).

Together these results indicated that the enhanced memory of *Wt1Δ* mice impacts the abilities of these mice to learn new experiences and flexibly modify their behavior to adapt toward changing environments. These results suggest that sequential learning would be impaired in *Wt1Δ* mice.

To obtain further experimentally testable predictions about possible effects of WT1 on sequential memory, we developed a toy control theory-based model of an information processing and response system. In the model we postulated that experience activates two parallel pathways: a memory-strengthening pathway that includes transcription factors like CREB and EGR1, and a memory-weakening pathway that includes transcription factors such as WT1. Together the pathways control the activity level of effectors to regulate balance that dictates the level of memory retention. *A priori.,* the total number of effectors could either be in excess of that needed to encode multiple experiences, or they could limit the encoding capacity of the cortico-hippocampal circuit (Fig.7a). We used the computational toy model to run simulations to study the effect of varying the activity of the memory weakening pathway for a fixed stimulus. The results of the simulations are shown in (Supplementary Fig. 7a). The model predicts that if the memory capacity of the cortico-hippocampal circuit is limiting, then over-representation of the first experience could interfere with the ability to acquire subsequent experiences. Alternatively, if effectors were not limiting, then reducing WT1 levels could enhance the ability to memorize both experiences (this is shown schematically in the bar graphs in Fig. 7a and in the simulation results in Supplementary Fig. 7b; refer to methods section Table S1:Model Parameters for a complete list of model parameters).

**Figure 7:**
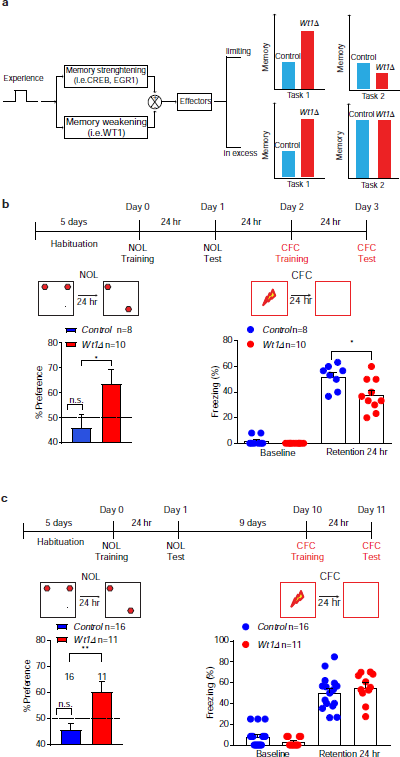
Consequences of WT1-mediated impaired memory flexibility, **a,** Proposed mechanism of WT1’s effect on memory regulation. An initial experience such as task 1 (NOL) activates both pro-memory strengthening and pro-memory weakening pathways. When the memory weakening pathways are inhibited by depletion of WT1, there is prolonged memory for task 1. Retention of task 1 memory may or may not interfere with the ability to remember a task 2(CFC) based on the availability of effectors (limiting vs in excess), **b,** Schematic representation for short-interval sequential training in mice (top panel). *Wt1Δ* animals showed increased time spent exploring the new location when first trained in NOL (left panel) and tested 24 hr after training (dashed line indicates 50% preference. Data are expressed as mean ± s.e.m.; unpaired t test: *p=0.0422; t=2.207, df= 16). 24 hr after being tested in NOL mice were trained on CFC (right panel) and *Wt1Δ* animals spent significantly less time freezing than their *Control* littermates when tested 24 hours after training (data are expressed as mean ± s.e.m.; unpaired t test: *p=0.0161; t=2.691, df=16). **c,** Schematic representation for long-interval sequential training (top panel). *Wt1*Δ animals showed increased time spent exploring the new location when first trained in NOL (left panel) and tested 24 hr after training (dashed line indicates 50% preference. Data are expressed as mean ± s.e.m.; unpaired t test: **p=0.0036; t=3.212, df=25). 9 days after being tested in NOL, *Wt1Δ.* mice were trained on CFC (right panel) and compared to *Control* littermates, they spent comparable amount of time freezing when tested 24 hours after training (data are expressed as mean ± s.e.m.; unpaired t test: p=0.3816; t=0.8906, df=25).

We therefore tested whether WT1 depletion, which enhances memory for one learning, would *interfere* with new learning in a sequential behavioral paradigm. We first trained mice in NOL, which does not yield LTM at 24 hour after training in control mice, but does so in *Wt1Δ* mice (whereas control mice shows memory retention at earlier time points, e.g. one hour after training; Fig. 2b). We then exposed the mice to a second learning experience, CFC, which normally does induce LTM (as shown in Fig. 2b). As depicted in Fig. 6b left panel, as expected, *Wt1Δ* mice had significant LTM retention for NOL at 24 hours after training, while control mice did not. However, when *Wt1Δ* mice that first underwent the NOL experience, were exposed one day later to CFC training, they showed significantly reduced LTM for CFC at 24 hours compared to control mice (Fig. 7b), indicating that the first experience learned in the absence of WT1 impacts subsequent learning. To determine the duration of this active learning interference, we tested the effect of extending the interval of time between NOL and CFC learning to 10 days. Consistent with previous experiments, *Wt1Δ* mice showed enhanced 24 hour retention for NOL (Fig. 7c). The 10 days delay between the two sequential experiences resulted in no difference between the two groups in LTM for CFC (Fig. 7c), indicating that the interference effect is a decaying function of the process induced by the first learning experience and it is temporarily limited. This suggests that strengthening memory by removing WT1 limits behavioral flexibility and that this effect is temporarily restricted.

## Discussion

Understanding the contribution of mechanisms of forgetting is critical for understanding memory storage and persistence. In this study, we identified a critical role of the transcriptional repressor WT1 in limiting memory strength, thereby promoting forgetting in order to offer flexibility to the learning and memory systems.

Since WT1, like activator transcription factors including C/EBP, cFos and Zif268 (Alberini 2009), is induced by LTP and learning, we conclude that the cascade of gene expression induced by learning and required for long-term memory requires specific transcriptional repression in addition to activation. Surprisingly, not many transcription factors have been studied in the context of forgetting and memory flexibility; one example is the transcription activator XBP1, which like WT1 is induced by learning ^67^, but which acts conversely as a positive regulator of hippocampal long-term memory and flexibility through transcriptional up-regulation of brain-derived neurotrophic factor (BDNF) ^68^. Given that the role of WT1 is to actively promote forgetting, transcription repression via specific DNA binding factors adds to other recently identified mechanisms of active forgetting (processes that counteracts memory consolidation and strengthening), which include neurogenesis and Rac1-, dopamine-, and Cdc42-mediated AMPA receptor endocytosis ^6^. Notably, the process of WT1-mediated active forgetting will occur via the function of its target genes, including several members of the retinoic acid signaling pathway (*Rdh5, Crabp2,* and *Ttr),* the immediate early genes *Arc* and *Fos,* as well as *Igf2* (and *Igfbp2),* which our data indicate to be involved in signaling through the IGF-2 receptor (IGF-2R). IGF-2R, also known as cation-independent mannose-6-phosphate receptor, binds IGF-2 (and other ligands) and targets it to lysosomal degradation. Hence, it is possible that lysosomal degradation serves to rebalance and complement the *de novo* protein synthesis and structural changes induced by learning. Of note, *Igf2* is one of the most well characterized WT1 target genes ^61^ and it has been previously shown to enhance synaptic plasticity, long-term memory and to prevent memory loss^63,69,^ mimicking some of our findings related to WT1 ablated animals. However, there are divergences as IGF-2 injected mice show enhanced fear extinction with similar memory flexibility^64,70,^ suggesting that induced IGF2 expression can explain only some of the behavioral effects observed in WT1-ablated animals (enhanced plasticity and long-term memory).

One additional observation is that WT1, by regulating its targets, might be involved in the regulation of homeostatic plasticity, which is the ability of glutamatergic neurons (among different neurons) to respond to periods of reduced or excessive activity by increasing or decreasing, respectively, their synaptic efficiency (synaptic scaling). Synaptic scaling occurs through enhancement or decrease of AMPA receptor-mediated excitatory synaptic transmission, which is in turn regulated by several molecular players ^71,72^. In this regard active forgetting has also been linked to cytoskeleton targeting mechanisms of synaptic remodeling^18,21,73^ and AMPA receptor recycling ^23,74^. The regulation mechanisms underlying homeostatic plasticity and AMPA receptor recycling are still only partially known, but they include some of the WT1 target genes, such as Arc ^75^, retinoic acid ^76,77^ and IGF-2 ^63^. Notably both retinoic acid and IGF-2 bind to the IGF-2 receptor, which regulates endocytosis, endosomal trafficking, and lysosomal degradation ^78^. We suggest that this regulation may influence AMPA receptor trafficking and levels, and therefore synaptic scaling, homeostasis, and plasticity.

Our results showing impaired subsequent learning after a first learning experience in the absence of WT1 suggest that the first learning is more strongly represented; hence, the mechanisms that counteract memory strengthening and consolidation normally are critical for memory formation. In fact, these mechanisms, by preventing over-consolidation of memories, allow learning flexibility that supports ability to adapt to changing conditions. Particularly important for pathologies in the area of trauma and anxiety was the observation that WT1-depleted mice trained in CFC show decreased extinction and an increased anxiety response, as measured by elevated plus maze. These behaviors are typical of anxiety disorders including PTSD, in which it is well known that memories of the aversive experience and traumas have been over-consolidated ^79^. As WT1 ablation in the hippocampus does not affect short-term memory, we suggest that the role of WT1 in forgetting is either to counteract memory consolidation or to impair retrieval, and further studies will be needed to understand this issue.

Accurate consolidation of long-term memories in the cortico-hippocampal circuit relies on coordinated activity in two major inputs, both originating in the entorhinal cortex but activating hippocampal CA1 neurons either directly, or through the trisynaptic pathway. At the level of CA1 neuron, a nonlinear response to synaptic input might underlie its capacity to function as a coincidence detector to decipher the synergistic effect of activating both pathways ^80^. We reported here that depletion of WT1 from the CA1 pyramidal neurons leads to a significant increase in excitability (Fig. 3f and 3h), to LTP enhancement (Fig. 3d and 3g), and to alteration of the entire intra-hippocampal circuit response (Fig. 4). Depletion of WT1 eliminated the ability of CA1 neurons to enable this circuit level computation, as the requirement for dual input to CA1 neurons is no longer needed to produce LTP.

Lastly, we speculate that the identification of WT1 as a new transcriptional regulator of memory persistence and memory flexibility may have potential implications for the treatment of those neurological conditions where memory is inflexible and excessively resistant to disruption, such as PTSD and OCD.

## Acknowledgements

The authors thank P.Y. Chuang for his guidance with the *Wt1Δ* mice colony and G. Jayaraman for her assistance with biochemical experiments. The authors want to thank Rachael L. Neve at MIT Viral Core for her assistance in generating the HSV viruses. This research was supported by NIH grants GM54508 and Systems Biology Center Grant GM071558 (PI: Iyengar), R37-MH065635 (PI: Alberini).

## Author Contributions

The project was designed by RI, CM, LM, RDB, CMA. CM conducted most of the biochemical experiments, the surgeries in rats and the behavior experiments in rats. LM conducted all the behavior experiments in mice. EG and JMS performed field recordings experiments while NT conducted patch clamp experiments. JH analyzed mRNAseq data and did enrichment analysis. SAS and VG helped with inhibitory avoidance and contextual fear conditioning experiments in rats. GEH and SJR provided equiμment and guidance for mice behavioral experiments. HA designed and assisted with the cloning strategy for WT1-HSV viruses. EUA helped with microscopy. VH provided the *Wt1*^*fl*/*fl*^ animals and provided expert insight in WT1 function and genetic manipulation. MB developed and ran the memory homeostasis model. CM, RDB, CMA and RI wrote the manuscript. All authors assisted in editing the manuscript.

## Methods

## Replication, blinding and statistical analysis

Experiments were run at least three separate times. For the *wt1Δ*-mRNA seq experiment, the results represent two different biological replicates. For behavior experiments the results are obtained from pulling together multiple animals from at least two different cohorts. Details of replicates are provided in each experiment. No statistical methods were used to predetermine sample sizes, but our sample sizes are similar to those reported in previous publications.

For all the electrophysiology and behavior experiments, the experimentalists were blind to the mice genotype or to the type of oligonucleotide or AAV/HSV virus treatment during the entire data gathering process. Only after the data were pooled and analyzed was the coding for the different groups revealed.

Unless otherwise stated, data are represented as mean ± s.e.m.. All the statistical analyses were run in GraphPad Prism 7.

## Data availability

The data that support the findings of this study are available from the corresponding authors upon request. Accession codes, unique identifiers and additional information on publicly available datasets are reported in the Reporting Summary file.

### Code availability

Custom written Matlab scripts were used for the control theory-based model of WT1 function. The code is available upon request.

### Research Animals

All experiments involving animals were performed according to ethical regulations and protocols approved by the internal Animal Care and Use Committee at Icahn School of Medicine at Mount Sinai.

### Cell fractionation and transcription factor activation arrays

Nuclear and cytosolic extracts were isolated according to standard procedures using low speed centrifugation. All buffers contained protease and phosphatase inhibitors. Tissue was lysed using a motorized Potter-Elvehjem homogenizer (∼10 strokes) in Buffer A (20 mM HEPES (pH 7.4), 40 mM NaCl, 3 mM MgCl, 0.5 % NP-40, 10 % glycerol, 1 mM DTT). Homogenized tissue was left for 10 min on ice, and lysates were spun at 500 g for 10 min at 4 °C to pellet nuclei. Nuclei were washed gently in Buffer B (20 mM HEPES (pH 7.4), 40 mM NaCl, 3 mM MgCl, 0.32 M Sucrose, 1 mM DTT) and spun at 500 g for 10 min at 4 °C. Nuclei were then resuspended using equal volumes of Buffer C (20 mM HEPES (pH 7.4), 40 mM NaCl, 1.5 mM MgCl, 25% glycerol, 1 mM DTT) and of Buffer D (20 mM HEPES (pH 7.4), 800 mM KCl, 1.5 mM MgCl, 1% NP-40, 25% Glycerol, 0.5 mM EGTA, 1 mM DTT). Samples were then rotated at 4 °C for 30 min to extract nuclear proteins and the resulting lysates were then spun at 13, 000 Rμm for 20 min at 4 °C.

The supernatant containing nuclear proteins was used to study transcription factors activation using the Panomics Combo Protein-DNA Array (Affymetrix, MA1215, now sold by Isogen Life Science). Each array membrane is spotted with 345 oligonucleotides that correspond to consensus binding sites for different transcription factors. The location on the array of each consensus binding site, as well as the complete protocol are available in the manufacturer’s website (http://www.isogen-lifescience.com/tf-protein-dna-array)). Five micrograms of nuclear extract were incubated with the biotinylated probe mix from the array kit for 30 min at 15°C. These probes are also transcription factor consensus binding sites that are complementary to the oligonucleotides spotted on the array. Probes that bound to transcription factors in the nuclear extract were purified by spin column separation, and bound probes were further purified from the transcription factors according to the manufacturer’s instructions. The purified probes were boiled for 3 min and hybridized overnight at 42°C to the array containing 345 oligonucleotide transcription factor consensus binding sites. The array was then washed, blocked, incubated with Streptavidin-HRP, and visualized by enhanced chemiluminescence. The blot was scanned and spot intensities were quantified using Image J.

For each condition (control and stimulated 30 minutes), ten CA1 regions were dissected from hippocampal slices obtained from at least three different animals and were pooled together in order to obtained sufficient nuclear extracts (5-10 *μg*). We compared extracts from unstimulated (control) slices with extracts from slices that were stimulated with Strong-HFS (see field recordings section within electrophysiology methods) and collected 30 minutes after stimulation.

### Gel Shift assay-EMSA

DNA probes were prepared by annealing complementary single-stranded oligonucleotides with 5’GATC overhangs (Genosys Biotechnologies, Inc.) and labeled by filling in with [α-32P]dGTP and [α-32P]dCTP using Klenow enzyme. For the CFC experiment, DNA probes were prepared using the LightShift Chemiluminescent EMSA Kit (Thermo Scientific) where complementary single-stranded transcription factor binding consensus sequence was first biotinylated using the Biotin 3’ End Labeling Kit (Thermo Scientific) and then annealed. In both experiments nuclear extracts were incubated with labeled DNA probes for 30 minutes at room temperature (22-24 °C). For the LTP experiment DNA-binding complexes were separated by electrophoresis on a 5% polyacrylamide-Tris/glycine-EDTA gel which was dried and exposed to X-ray film. For the CFC experiment protein/DNA complexes were separated using a 6% DNA retardation gel (Invitrogen)that was electroblotted into a Biodyne B membrane (Thermo Scientific), incubated with Streptavidin-HRP (Thermo Scientific) and visualized by ECL according to the manufacturer’s instructions. The consensus sequence used for WT1 was: 5’-AATTCGGGGGCGGGGGCGGGGGCGGGGGAGGGGCGC-3’ and its complementary sequence. For the CFC experiment binding was confirmed using an additional consensus sequence 5’-TCCTCCTCCTCCTCTCCC-3’.

For the LTP experiment slices were stimulated using Strong-HFS protocol (see field recordings section within electrophysiology methods); for the CFC experiment, animals were trained using three footshocks (2-sec each, 0.65 mA, 1 min apart). The control animals (indicated as “C”) remained in the conditioning chamber for the same amount of time as the ones receiving the shock (indicated as “S”) but without receiving any footshock.

### Real time quantitative RT-PCR

Hippocampal total RNA was extracted with TRIzol (Invitrogen) and 1 μg of total RNA was reverse-transcribed using SuperScript III First-Strand Synthesis System (Invitrogen, ThermoScientific, catalogue #18080-051). Real-time PCR was performed using 7500RT PCR System (Applied Biosystems). 1 μl of the first-strand cDNA was subjected to PCR amplification using a QuantiTect SYBR Green PCR kit (Qiagen). IGF-II primers (forward: 5’-CCCAGCGAGACTCTGTGCGGA-3’; reverse, 5 ‘-GGAAGTACGGCCTGAGAGGTA-3’);Forty cycles of PCR amplification were performed as follows: denaturation at 5°c for 30s and extension for 30s at 72°C. GAPDH (forward, 5 -TGCACCACCAACTGCTTAGC -3’; reverse, 5 ‘-GGCATGGACTGTGGTCATGA -3’) was used as internal control. To determine the relative quantification of gene expression the cycle threshold method (*C*_T_) was used.

### Immunohistochemistry

Rats and mice were deeply anesthetized, perfused using 4% paraformaldehyde and coronal or hippocampal brain sections were obtained using a vibratome (Leica VT 1000S vibratome; 40 μm) or a cryostat (Leica CM1850; 15μm). Brain slices were then blocked with 3% normal goat serum (Vector), 0.3% Triton X-100 (Sigma-Aldrich), 1% BSA (Sigma-Aldrich) for 2 hr at room temperature and incubated with the appropriate primary antibody: rabbit monoclonal WT1 (for staining in Figure 3 a Santa Cruz, catalogue #SC-192 (C-19); for staining in Figure 3b Novus Biological, catalogue #NBP1-40787; for staining in Figure 3c Abcam, catalogue #ab52933); mouse monoclonal glial fibrillary acidic protein, GFAP (Cell Signaling, catalog #3670); mouse monoclonal β-tubulin (Cell Signaling, catalog #86298). An antibody against green fluorescent protein-GFP (chicken anti-GFP, from Aves Labs Inc., catalogue #GFP-1020) was used to check viral spread in WT1 over-expression experiments using AAV and HSV viruses (Fig. 2c and 2d). After incubation with primary antibodies, sections were washed and incubated with secondary antibodies complexed to either Alexa Fluor 568 or Alexa Fluor 488 dyes (Invitrogen, ThermoFisher). After washing, Hoechst 33342 (Invitrogen) was used to label nuclei. Sections were then mounted and imaged using a confocal microscope (Zeiss LSM 800).

### Western Blotting

We used either CA1 regions dissected from 400 μm-thickness hippocampal slices or dorsal hippocampi, homogenized in proportional volumes of ice-cold lysis buffer using a motorized Potter-Elvehjem homogenizer (∼10 strokes). The lysis buffer consisted of 50 mM Tris-HCl, pH 7.4, 100 mM NaCl, 1 mM EDTA, 0.5% Sodium Deoxycholate, 1% NP-40, 0.1% SDS, 0.5 mM μmSF, 1 μM Mycrocystine, 1ug/ml Benzamidine, 2 mM dichlorodiphenyltrichloroethane (DTT), 1 mM Sodium orthovanadate, 2 mM Sodium Fluoride, 1 mM EGTA; protease inhibitor cocktail (Sigma-Aldrich) and phosphatase inhibitor cocktails 2 and 3 (Sigma-Aldrich) were added according to manufacturer’s instructions. Lysates were cleared by centrifugation at 14,000 Rμm for 10 min. Protein concentration was determined using Bradford reagent (Biorad). 20-50 μg of total protein was loaded per well, into 10% SDS-PAGE and transferred to supported nitrocellulose membranes (pore size 0.2 μm, Biorad), followed by western blotting and chemiluminescence detection. The following antibodies were used: rabbit polyclonal WT1 (custom-made, against a synthetic rat-specific peptide; GeneScript), rabbit monoclonal WT1 (Novus Biological, catalogue #NBP1-40787), mouse monoclonal β-tubulin (Cell Signaling, catalog #86298), mouse monoclonal β-actin (Sigma-Aldrich, catalogue #A4700). Films were scanned, and net intensity was analyzed using ImageJ.

For the experiments in Figure 1e-g the time point considered was 30 min either after Strong-HFS delivery or behavioral training; total protein lysates were collected from CA1 regions (Figure 1e) or from dorsal hippocampi (Figure 1f and 1g).

For the LTP experiment slices were stimulated using Strong-HFS protocol (see field recordings section within electrophysiology methods); for the CFC experiment, shocked animals received 3 mild footshocks (2-sec each, 0.65 mA, 1 min apart) during training session. The un-shocked animals (Context only, Ctx only) remained in the conditioning chamber for the same amount of time as the shocked ones without receiving any footshock. For the IA experiment trained animals (Trained 30 min) were compared to Naive ones.

### Cannulae implants and hippocampal injections of ODNs or HSV viruses in rats

Rats were cannulated as previously described ^1^. Rats were anesthetized with a solution containg a mix of ketamine (70 mg/and xylazine (10 mg/kg, intraperitoneal), and a stainless-steel guide cannulae (22 gauge) were stereotactically implanted to bilaterally target the dorsal hippocampus (4.0 mm posterior to the bregma, 2.6 mm lateral from midline and 2.0 mm ventral) ^2^. The rats were returned to their home cages and allowed to recover from surgery for 7 to 10 d.

### WT1 knock-down

All hippocampal injections consisted of 2 nmol in 1 μl per side (unless otherwise specified) of either WT1 antisense oligodeoxynucleotide combo (WT1-AS= 1 nmol of WT1 antisense 1 + 1 nmol of WT1 antisense 2) or scrambled oligodeoxynucleotide combo (SC-ODN= 1 nmol Scrambled 1 + 1 nmol Scrambled 2) both diluted in phosphate-buffered saline (PBS) at pH 7.4. The sequences used were the following: WT1 antisense 1: TCGGAACCCATGAGGTGCGG; WT1 antisense 2: TCGGAACCCATGGGGTGC; Scrambled 1:GGTGGTAGAACGCCGTACCG; Scrambled 2: GGTGGTAGAACGCCGTCC. The scrambled oligonucleotides, which served as a control, were designed to lack homology to any rat sequence in GenBank, and contained the same base composition but in a randomized order. Both antisense and scrambled oligonucleotides were phosphorothioated on the three terminal bases of both 5’ and 3’ ends to increase their stability and were reverse phase purified (GeneLink).

For electrophysiology experiments, male Sprague-Dawley rats were used. Animals received a single injection of oligonucleotides 2 hr before being sacrificed, and their brains were dissected (see Fig. 3d for schedule diagram). For electrophysiology experiments one side of the brain was always injected with WT1-AS and the other side of the brain with SC-ODN. For all the behavior experiments either male Sprague-Dawley or Long-Evans rats were used and no differences between the strains were observed. For behavior experiments, animals received two injections of oligodeoxynucleotides 2 hr apart and 2 hr before training (see Fig. 2a for schedule diagram); animals were injected bilaterally with either WT1-AS or SC-ODN.

### WT1 over-expression

Herpes-simplex Virus (HSV): for non-conditioned over-expression of WT1 via HSV we used a p1005 based HSV vector co-expressing GFP and WT1-IsoformD (WT1-HSV). In this system, GFP expression is driven by a cytomegalovirus (CMV) promoter, while the WT1-isoformD is driven by the IEF4/5 promoter. HSV virus expressing GFP alone was used as a control (CTR-HSV). We injected 2 μl of HSV vectors in each hemispheres (titer 0.5x10^^^9 infectious unit/ml, Virovek, Hayward, CA) using a 28-gauge needle that extended 1.5 mm beyond the tip of the guide cannula and connected via polyethylene tubing to a Hamilton syringe. The infusions of HSV viruses were delivered at a rate of 0.33 μl min^-1^ using an infusion pump (Harvard Apparatus). The injection needle was left in place for 10 minutes after the injection to allow complete diffusion of the solution. Rats were randomized to different treatments.

Over expression via AAV: For non-conditioned WT1 expression via AAV, we used AAV8.2-EF1a-WT1-PP2A-GFP (WT1-AAV) and AAV8.2-EF1a-PP2A-GFP (CTR-AAV; both vectors were 1 × 10^13^ vg/ml, Virovek, Hayward, CA) as a control. AAV vectors were injected using a 33 Ga needle attached to a 5ul syringe (Hamilton) 2 μl in each hemisphere over a 10 min period.The needle was left in place for 10 min to allow for efficient diffusion before removal. Rats were randomized to different treatments.

To verify proper placement of cannula implants or viral injection, rats were sacrificed at the end of the behavioral experiments, their brains removed and fixed with 10% Δsections (40 μm) and the hippocampus region was examined under a light microscope (for cannulae placement) or confocal microscope (for viral injection). Animals where cannulae were misplaced, viral expression was mostly spread outside of the hippocampus, and serious tissue damage was observed were excluded from the experimental groups.

## Generation of functionally deficient WT1 mice

Forebrain-specific deletion of *Wt1* was achieved by crossing animals homozygous for the conditional *Wt1* knockout allele (*Wt1*^fl/fl^) ^3^ with a transgenic line, *Camk2a-Cre*, (B6.Cg-Tg(Camk2a-cre)T29-1Stl/J; Jackson Lab: (http://jaxmice.jax.org/strain/005359.html) in which Cre recombinase expression is driven by the 7.8kb promoter of Ca2+/calmodulin-dependent protein kinase II alpha subunit ^4^ Progeny were crossed to obtain *Wt1* ^*fl*/*fl*^; *Camk2a-Cre* (referred through the paper as *Wt1Δ* mice) and littermate control animals (referred through the paper as *Control* mice). Expression of Cre recombinase resulted in the in-frame deletion of exons 8 and 9 [see Figure 1e of Gao *et al*.], and generated a truncated allele encoding a shortened non functional WT1 protein lacking zinc fingers 2 and 3. Expression of the recombined Wt1 allele was detectable in the mouse forebrain (Fig. Supplementary 3a), and its detection was performed as previously described using ^3^ the following primers: Primer WT1 Delta Forward 5’ GCT AAC ATA TGG GAG ACA TT 3’ and Primer WT1 Delta Reverse 5’ TGC CTA CCC AAT GCTCAT TG 3’. As reported by others, heterozygous Wt1 mice develop kidney nephropathy and glomerulosclerosis ^5^, which we have not observed at any time in the *Wt1Δ* mice. To further address this issue, we evaluated proteinuria since loss of kidney function is associated with increased levels of proteins in the urine. Using Chemstrips (Roche), we found that there was no significant difference between proteinuria levels of *Wt1Δ* mice compared to their control littermates as indicated by the color of the top strips (Fig. Supplementary 3c). We further confirmed that kidney function was normal and that there was no significant difference in the enzymatic values of *Wt1Δ* mice through a pathology screening of their blood samples performed at the Comparative Pathology Center of Mount Sinai (Fig. Supplementary 3d).

To genotype the animals we used the following primers for the LoxP allele: Primer LoxP Forward 5’ CCT TTT ACT TGG ACC GTT TG 3’ and Primer LoxP Reverse 5’ GGG GAG CCT GTT AGG GTA 3’. For the Cre allele we used the following primers: Cre Primer Forward 5’ GCG GTC TGG CAG TAA AAA CTA TC 3’ and Cre Primer Reverse 5’ GTG AAA CAG CAT TGC TGT CAC TT 3’ (as indicated in the genotyping section by Jackson lab at (http://jaxmice.jax.org/strain/005359.html).

*Wt1Δ* animals were viable and had a normal life span, normal body weight, normal fertility and a normal growth rate compared to control littermates.

Throughout the study control wild-type littermates are indicated as *Control* and they comprise the following subgroups: *Wt1*^+/+^; *Camk2a-Cre* positive, *Wt1*^+/+^; *Camk2a-Cre*negative, *Wt1*^*fl*/^+ *Camk2a-Cre* negative, *Wt1*^*fl*/*fl*^; *Camk2a-Cre* negative. These were grouped together for both electrophysiology and behavior experiments, since they were no statistically different between the genotypes.

## Hematoxylin and eosin (H & E) staining

H & E staining was performed in order to verify if there was any macroscopic abnormality in brain tissue of *Control* and *WT1Δ* mice. Animals were deeply anesthetized with a solution containing ketamine + xylazine and perfused transcardially with ice-cold 10% formalin. The brains were embeded in paraffin and sliced into 2 μm thick sections for staining. The sections were de-paraffinized in xylene, rehydrated in graded ethanol series, stained with Mayer’s Haemalaun (Carl Roth, Karlsruhe, Germany) for 5 min, washed again, and stained with 1% eosin (Carl Roth, Karlsruhe, Germany for 2 minutes). The sections were washed in water, dehydrated in graded ethanol series, treated with xylene and mounted for imaging (Fig. Supplementary 3b).

## Behavioral Assays Contextual Fear Conditioning and Extinction

Mice or rats were handled for 3 minutes per day for 5 days before training. The conditioning chamber consisted of a rectangular Perspex box (VFC-008: 30.5 × 24.1 × 21.0 cm, Med Associates) with a metal grid floor (Model ENV-008 Med Associates) through which a footshock was delivered. The experiment was conducted in a sound-attenuated room, with low levels of light and white noise background. Unless otherwise specified, during training session, animals were allowed to explore the box for 2 minutes prior the delivery of a single footshock (2-sec, 0.65 mA); after that the animals remained in the chamber for two additional minutes before returning to their home cages. They were tested 24 hours after the training and 30 days after training (mice only). Test session consisted in placing the animals back into the conditioning chamber for 5 min in the absence of any footshock. For the mice memory extinction experiment, after CFC training, animals were placed into the conditioning chamber for 5 consecutive days, 5 min each day in the absence any footshock and freezing was scored. Sessions were recorded using a digital video camera, and freezing behavior defined as lack of movement besides heart beat and respiration, was scored every 10 seconds by trained observers blind to the experimental conditions. The number of scores indicating freezing were calculated as a percentage of the total number of observations ^6^. For experiments with WT1 over-expression freezing was scored using Ethovision (Noldus Information Technology).

## Inhibitory Avoidance (IA)

IA was carried out as described previously^7^.The IA chamber (Med Associates) consisted of a rectangular Perspex box divided into a safe compartment and a shock compartment.. The safe compartment was white and illuminated, whereas the shock compartment was black and dark. The chamber was located in a sound-attenuated, non-illuminated room. Foot shocks were delivered though the grid floor of the shock chamber via a constant current scrambler circuit. During training sessions, each rat was placed in the safe compartment with its head facing away from the door. After 10s the door separating the compartments was automatically opened, allowing the rat access to the shock compartment; the rats usually enter the shock (dark) compartment within 10–20 s of the door opening. As soon as rats stepped into the shock compartment a mild footshock was delivered. (0.60 mA for 2 s). For the western blot experiment (Fig. 1g) using IA extracts, animals were euthanized 30 minutes after training using halothane and their brains dissected. Dorsal hippocampi from trained animals were compared to dorsal hippocampi obtained from naive controls (animals that remained in their home cages).

## Novel Object Location (NOL)

For both mice and rats experiments, animals were allowed to familiarize with the arena for 5min each day for 3 consecutive days before training. The arena consisted of a box (44.4cm × 44.4cm × 31.5 cm for rats and 28cm × 28cm × 20cm for mice) with one of the walls covered with dark/opaque paper, while the other three contained visual cues. Arena was placed in a room with a low level of light and sound-proof. During the training session two identical objects (Lego®) were placed into the arena. Animals were placed in the middle of the arena always facing the wall covered with the dark/opaque paper and were allowed to freely explore the objects for 10 minutes, returning to their home cages afterwards. During testing animals were placed back into the arena for 5 minutes and one object was moved to a different location. Object location was counterbalanced during training and test. Object exploration was defined as the orientation of the animal’s nose towards the object at a distance ≤ 2 cm or as the animal placing its forepaws on the object; climbing on the object was not considered exploration. The objects and the arena were cleaned with 70% ethanol between animals to avoid olfactory cues. For NOL experiments with rats the sessions were videotaped and scored by an experimenter blind to experimental conditions; for NOL experiments with mice the sessions were scored using Ethovision (Noldus Information Technology). Memory retention was measured as % Preference calculated as the time spent exploring the object in the new location (N) relative to the total exploration time (N + familiar (F)) (% Preference = (N / (N+F)*100) ^8^.

## Open Field

For the locomotion experiment in rats, animals were allowed to freely explore for 5 min an open field arena (44.4cm × 44.4cm × 31.5 cm) divided into 16 imaginary quadrants. Locomotion was calculated as total number of crossings in the open field. An observer blind to experimental procedures scored the experiments.

For the mice locomotion experiment, animals were allowed to explore an empty arena (34cm × 34cm × 23 cm) for 10 minutes during which the total distance traveled as well as the time spent in the center or periphery of the arena were recorded using a videotracking system (Ethovision, Noldus Information Technology).

## Spontaneous Alternation and Reversal Learning in a Y maze

Spontaneous alternation and reversal learning were performed as described previously ^9^. The Y-maze consisted of three white opaque arms (Med Associates) with sliding doors at the entrance of each arm. During spontaneous alternation test animals were allowed to freely explore the three arms from the center of the maze for 10 minutes and spontaneous alternation was defined as successive entries into each of the arms on overlapping triplets sets (e.g. ABC, BCA, CAB, etc). The percentage of alternation was calculated by as the ratio of total alternations to possible alternation (total arm entries -2) × 100.

For the reversal learning experiment mice were single housed, food restricted and monitored daily until they reached 85% of their original weight before starting the experiment and during testing. They were given 1/2 food pellet (LabDiet 5053) and 1 fruit loop (Kellog’s) each day. The habituation phase was identical to spontaneous alternation. During the acquisition phase, one arm of the maze was chosen as the “correct arm” and baited with half of a fruit loop. The animals were initially restrained in the “start arm” for one minute and then allowed to explore between the two arms. The acquisition phase consisted of 10 consecutive trials per day for 2 days (each day divided in 2 blocks of 5 trials each). Memory was calculated as the percentage of correct choice over each block of trials. During the reversal learning phase the “correct arm” was switched. The “correct arm” was counterbalanced between animals.

Both experiments were scored by an observer blind to the experimental conditions.

## Marble Burying Test

Marble burying test was carried out as previously described ^10^. Regular rat cages were used and filled with approximately 5 cm deep bedding tamped down to make a flat, even surface. A regular pattern of 20 glass marbles was positioned on the surface of the bedding, spaced regularly, about 4 cm apart one from the other. Each animal was left in the cage for 30 minutes and the % marbles buried was calculated as the number of marbles buried to approximately 2/3 of their depth over the total number of marbles X 100.

## Elevated Plus Maze

The elevated plus maze consisted of black Plexiglas fitted with white bottom surfaces to provide contrast and was placed 60 cm above the floor. The four arms (2 open and 2 closed) were interconnected by a central platform. Test was conducted as previously described ^11^. Briefly, mice were placed at the center of the maze were allowed to freely explore it for 5 min under red-lighting conditions. Using a videotracking software (Ethovision, Noldus Information Technology) and a ceiling-mounted camera the time spent in the open and closed arms as well as the number of entries in the closed and open arms was recorded and further analyzed.

## Plantar test (Hargreaves method)

To assess mice nociceptive response, animals were placed in a clear plastic chamber (45 cm × 40 cm, divided in 12 small animal enclosures, IITC Life Science) with a glass floor and allowed to acclimatize to the room and to the apparatus for 2 hours. After the acclimation period, the radiant heat source (infrared beam) was positioned under the glass floor directly beneath one of the animal’s hind paws. The radiant heat source creates a 4 × 6 mm intense spot on the paw. The paw withdrawal latency was determined using an electronic stopwatch coupled to the infrared source that switches off when the animal feels discomfort and withdraws its paw; a cutoff of 20 sec for paw withdrawal was set up.

## Electrophysiology

### Field recording

Male Sprague-Dawley rats (6-8 weeks old) or approximately 3 months old mice (either *Control* or *Wt1Δ* mice) were deeply anesthetized with isoflurane and decapitated. The brain was rapidly removed and chilled in ice-cold artificial Cerebro Spinal Fluid (ACSF) containing (in mM) 118 NaCl, 3.5 KCl, 2.5 CaCh, 1.3 MgSO4, 1.25 NaH2PO4, 24 NHCO3, and 15 glucose, bubbled with 95% O_2_/5% CO_2_. Transverse slices of dorsal hippocampus (400 μm thick) were made on a tissue chopper at 4°C, and then placed in an interface chamber (ACSF and humidified 95% O_2_/5% CO_2_ atmosphere), where they were maintained at room temperature for at least 2 h. For recording, slices were transferred to a submersion chamber and superfused with ACSF at 31 ± 1°C. Monophasic, constant-current stimuli (100 μs) were delivered with a bipolar stainless steel electrode positioned in stratum radiatum of area CA3, and field EPSPs (fEPSPs) were recorded in stratum radiatum of area CA1, using electrodes filled with ACSF (Re = 2-4 MΩ). For all slices, initial spike threshold exceeded 2 mV. Signals were low-pass filtered at 3 kHz and digitized at 20 kHz, and analyzed using pClamp 9 (Molecular Devices). Two HFS protocols were used: Weak-HFS, consisting of two trains separated by 20 s, each consisting of 100 stimuli delivered at 100 Hz at an intensity that initially evoked a fEPSP measuring 20% of spike threshold; and Strong-HFS, identical to Weak-HFS but delivered at an intensity that initially evoked a fEPSP of 75-80% of spike threshold. In all experiments, the stimulation protocol was delivered at least 30 min after transfer of the slices to the recording chamber, when the basal fEPSP had been stable for at least 20 min. Control slices were placed in the recording chamber and subjected only to test stimuli (0.033 Hz). Drug preincubations, when used, were performed at room temperature in submersion maintenance chambers containing ACSF saturated with bubbling 95% O_2_/5% CO_2_. Drugs were prepared as stock solutions and diluted to final concentrations in ACSF before use.

In slices where both the TA→CA1 and SC→CA1 inputs were activated, stimulating electrodes were placed both in proximal stratum radiatum near the CA1/CA2 border (to activate Schaffer collaterals) and in the lacunosum-moleculare within CA1 (to activate the perforant path). For the baseline period, slices were stimulated every 30 s, alternating between Schaffer collaterals and perforant path. The perforant path was activated with theta-burst stimulation (TBS) consisting of 10 bursts at 5 Hz, 4 pulses per burst at 100 Hz, using 250 μA stimuli. The Schaffer collaterals were stimulated with the same TBS pattern, delayed 20 ms delay relative to the perforant path, at an intensity that initially evoked 90% of the spike threshold. Recording electrodes were positioned in stratum radiatum and stratum lacunosum-moleculare. All slices had a spike threshold of at least 1.8mV in stratum radiatum.

For recordings in the presence of bicuculline, the brain was rapidly removed and chilled in ice-cold ACSF containing (in mM) 118 NaCl, 2.5 KCl, 4 CaCh, 4 MgSO4, 1.25 NH2PO4, 24 NaHCO_3_, and 15 glucose, bubbled with 95% O_2_/5% CO_2_. Transverse slices of dorsal hippocampus (400 μm thick) were made on a tissue chopper at 4°C, and then placed in an interface chamber (ACSF and humidified 95% O_2_/5% CO_2_ atmosphere), where they were maintained at room temperature for at least 1 h. The CA3 region was then dissected from CA1 region and slices were placed in a submersion chamber for 0.5-2.5 h before being transferred to the recording chamber. A Weak-HFS was delivered at a stimulus strength that evoked a fEPSP measuring 25-30% of spike threshold in bicuculline. All other conditions were as described above. Bicuculline was suspended in water to 10 mM and diluted to 10 μ? in ACSF immediately before the experiment began.

### Whole-cell recording

Adult male Sprague-Dawley rats (250-300 g) were deeply anesthetized with isoflurane and transcardially perfused with ice-cold ACSF. For experiments on excitability (Figure 3f), the ACSF contained (in mM): NaCl (128), D-glucose (10), NaH2PO4 (1.25), NaHCO3 (25), CaCl2 (2), MgSO4 (2), and KCl (3), bubbled with 5% CO2 / 95% O2 (pH=7.3, 290-300 mOsM). Following perfusion, the brain was rapidly removed and chilled in ice-cold sucrose-ACSF containing (in mM): sucrose (254), D-glucose (10), NaH_2_PO4 (1.25), NaHCO_3_ (25), CaCl_2_ (2), MgSO_4_ (2), and KCl (3) (pH=7.3, 290-310 mOsM). Coronal slices of dorsal hippocampus (200 μm thick) were prepared using a vibratome in ice-cold sucrose-ACSF, and were allowed to recover submerged in bubbled ACSF for 45 minutes at 33 ± 1° C, and thereafter at room temperature. Slices were transferred to a submersion recording chamber and perfused with ACSF (2 mL/min) at room temperature. CA1 pyramidal neurons were identified using IR DIC optics, and whole-cell recordings were obtained with an Axopatch 1D amplifier. Signals were low-pass filtered at 2 kHz and digitized at 20 kHz, and no adjustment was made for pipette junction potential. Membrane excitability was tested in current clamp mode using pipettes containing (in mM): K gluconate (115), KCl (20), MgCl_2_ (1.5), phosphocreatine-Tris (10), Mg-ATP (2), Na-GTP (0.5), and Hepes (10) (pH=7.3, 280-285 mOsM; 3.5-4.5 ?Ω). The membrane was depolarized with a series of ten 200 ms-long current steps, increasing from 10 to 100 pA from a holding potential of -70 mV.

For recording spontaneous and miniature EPSCs (mEPSCs) (Supplementary Fig. 6a and 6b), slice preparation and recordings were performed in modified ACSF containing (in mM): NaCl (128), D-glucose (10), NH2PO4 (1.25), NHCO3 (25), CaCh (2), MgCh (2), and KCl (3) (pH=7.3, 290-300 mOsM), using pipettes filled with (in mM): Cs-methanesulfonate (130), HEPES (10), EGTA (0.5), NaCl (8), TEA-Cl (5), Mg-ATP (4), Na-GTP (0.4), Na-phosphocreatine (10), and N-ethyl lidocaine (1) (pH=7.3, 280-285 mOsM; 3.0-4.5 ?Ω). mEPSCs were recorded in the presence of D,L-2-amino-5-phosphonovaleric acid (APV; 50 μM), gabazine (5 μM), and tetrodotoxin (0.5 μM). Spontaneous events were recorded in the absence of inhibitors. 3-5 minutes after breakthrough, gap-free recordings were obtained for 10 minutes. Only cells with stable input resistances (< 20% change as measured before and after the gap-free period) were included in the analysis. Template-based event detection was performed using Clampfit 10.3 (Molecular Devices). Templates were generated by averaging 5-10 events for each file, and the automated search results were verified manually.

## Molecules and Inhibitors used in electrophysiological experiments

Bicuculline was purchased from Tocris (catalogue #2503) and resuspended in ACSF to reach a final concentration used 10 μM. The antibody against the IGF2 Receptor (IGF2-R Ab) was purchased from R&D solutions (catalogue #AF2447) and used at a final concentration of 5 μMg/ml.

## Transcriptomic profiling by mRNA seq

For the mRNA seq experiments, total RNA was extracted using Trizol (Thermo Fisher) from CA1 regions isolated from rat hippocampal slices (Control vs LTP 90 minutes). A pool of approximately 10 CA1 regions collected from at least 3 different animals were necessary in order to obtain approximately 1 μMg of total RNA for each condition. For the experiment relative to Wt1Δ mice versus wild type littermates, dorsal hippocampi from naïve untrained animals were used. For the experiment relative to acute WT1 knock-down in rats (WT1-ODN vs Scrambled-ODN, naïve untrained animals), dorsal hippocampus tissue surrounding the injection site was used. For all the mRNA sequencing experiments RNA integrity was checked by either the Agilent 2100 Bioanalyzer using the RNA 6000 Nano assay (Agilent, CA, USA). All processed total RNA samples had RIN value ≥9. The seq library was prepared with the standard TruSeq RNA Sample Prep Kit v2 protocol (Illumina, CA, USA). Briefly, total RNA was poly-A-selected and then fragmented. The cDNA was synthesized using random hexamers, end-repaired and ligated with appropriate adaptors for seq. The library then underwent size selection and purification using AMPure XP beads (Beckman Coulter, CA, USA). The appropriate Illumina-recommended 6 bp barcode bases are introduced at one end of the adaptors during PCR amplification step. The size and concentration of the RNAseq libraries was measured by the Agilent 2100 Bioanalyzer using the DNA 1000 assay (Agilent, CA, USA) before loading onto the sequencer. The mRNA libraries were sequenced on the Illumina HiSeq 2000 System with 100 nucleotide single-end reads, according to the standard manufacturer’s protocol (Illumina, CA, USA).

For the RNA-Seq data analysis Tophat 2.0.13 ^12^, bowtie 2.1.0 ^13^, samtool 0.1.7 ^14^ and cufflinks1.3.0 ^15^ were used. The rn5-bowtie2 index was generated with the command ‘bowtie2-build rn5.fa rn5’. The ‘rn5.fa’-file was downloaded from the UCSC genome browser. The mm10-bowtie2 index was downloaded from (http://bowtie-bio.sourceforge.net/bowtie2/manual.shtml). RefSeq geneTracks and GTF-files for the rn5 and mm 10 genome assembly were downloaded from UCSC genome browser. Common gene ids in the GTF-files were matched to individual transcript_ids using the corresponding official symbols obtained from the geneTracks files.

The likelihood to detected a lowly to moderately expressed gene in a particular sample depends on the total number of sequenced reads, especially in case of lower reads counts (<30, 000, 000) ^16^. Therefore it could happen that more genes are detected in a sample with a higher read count than in a sample with a lower read count. This experimental artifact might distort normalization including total reads normalization as well as upper quartile normalization that is applied in this study. Both normalization methods only change the number of reads that are associated with a gene, but not the number of identified genes. In consequence, the same number of reads might be distributed over a different number of (by chance) experimentally identified genes in two samples, introducing gene expression differences between the two samples that do not exist. To prevent such experimental artifacts reads we applied an additional computational step before read alignment and differentially expressed genes detection. Under the assumption that during the seq process every fragment has the same chance to be sequenced, we ensured that each sample had the same number of total read counts by randomly removing reads from those samples with higher read counts than the minimum read count.

Reads were aligned to the rn5 or mm10 genome using Tophat with the option ‘--no-novel-juncs’ and the refSeq-GTF-file (the option ‘csolexa1.3-quals’ was additionally chosen in case of the rat samples). Differentially expressed genes were identified using Cuffdiff with the options ‘--upper-quartile-norm’, ‘--frag-bias-correct’ against the rn5 genome and ‘--multi-read-correct’ and the refSeq-GTF-file.

In each analysis all differentially expressed genes (DEGs) that were statistically significant (FDR = 5%) were considered. DEGs with a minimum fold change of log_2_((FPKMcondition_1_+1)/(FPKMcondition_2_+1)) >= +/-log_2_(1.3) were submitted to pathway enrichment analysis as described below.

## Analysis of transcriptomic data

Enrichment analysis using mRNA seq data was performed similarly as previously described ^17^. The “Transfac_and_jaspar_pwms” library was downloaded from the EnrichR website^18, 19^ All human transcription factor gene associations were kept. Human target genes and transcription factors were replaced by their rat homologues based on the mouse informatics database (Mouse Genome Informatics [MGI], (http://www.informatics.jax.org, 5/24/2013) and the National Center for Biotechnology Information homologene database(http://www.ncbi.nlm.nih.gov/homologene/, 06/01/2018). Mouse gene-transcription factor associates were removed from the database.

To increase the statistical accuracy we removed all gene symbols in both databases that are not part of the RefSeq rn5 gene annotation and therefore could not be identified as differentially expressed. Similarly, we removed all differentially expressed genes that were not part of the “Transfac_and_jaspar_pwms” library. Right tailed fisher’s exact test was used for enrichment analysis and the negative logarithms to the basis 10 of the p-values were calculated.

## Control theory-based toy model of WT1 function

Input of an experience to the hippocampus is represented as a rectangular pulse. Neuronal activity in the hippocampus converts this pulse into a more long lasting output with respect to the time scale of the experiments (days), which we represent as a time integrator. Thus, the area under the rectangular pulse becomes a step function as inputs to memory-strengthening and memory-weakening pathways. We are unaware of any experimental data to suggest reasonable values for the magnitude of this step input, *u*, hence arbitrary values were chosen and *u* was subsequently varied to make a range of predictions (Supplementary Fig. 7). We model memory-strengthening and memory-weakening signaling as two first order processes in parallel. A first order process is governed by the following equation:

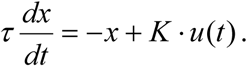

Here, *τ* is the time constant, *K* is the steady state gain, *u* is the input strength, *t* is time, and × is the dependent variable (in this case memory-strengthening or memory-weakening signal strength). We denote memory-strengthening with the subscript 1 and memory-weakening with the subscript 2. Because activation of one cell’s signaling could affect other non-activated cells, we take both gains (*K_1_* and K_2_), to be 3, reflecting signal amplification. However, in the model the effects of these gains and the input magnitude are indistinguishable, so our parameter variation exercise effectively explored both of these avenues. Additionally, we estimated from electrophysiological data that lack of functional WT1 induces an approximately 2.4-fold increase in the input signal strength, so in the case of *Wt1Δ* mice, we take the gains as 7.2. The time constants *τ*_1_ and *τ*_2_ were tuned to be consistent with the data in Figs. 1-3. Thus, τ _*1*_ for memory-strengthening signaling was taken as fast (0.5 hrs) and not affected by lack of functional WT1, whereas *τ* _2_ for memory-weakening signaling was taken as slow (36 hrs) and took a different value for *Wt1Δ* animals (144 hrs). These model parameters are summarized in the below Table S1:

The difference of these two process outputs was passed through a saturation function (based on neurobiological reasoning presented in the main text), to be fixed between 0 and 1, which we call “Pathway Activity”. Thus,

*Pathway Activity = sat*(*x*_1_-*x*_2_, 0, 1)

This “Pathway Activity” variable coarsely represents an amalgamated capacity for learning new events in the short-term. Based on the assumption of a finite amount of downstream effectors that interpret pathway activity, we define

*Effectors Available = 1–Pathway Activity.*

We specify that “*Memory^1^*” is a function of pathway and effectors dynamics by the following logic. In the absence of any past event, we can calculate the peak of *Pathway Activity* elicited by a particular event. This peak value is taken as the amount of capacity required to fully learn, which we call “*need*”. Then, we can calculate the *Effectors Available* elicited by a particular event as a function of time, given that other events may have already occurred previously, which we call “*have*”. *Memory* at each time point is defined as the *Pathway Activity* attributable to a particular event, divided by its maximum value, but weighted by the fraction *have/need.* Specifically,

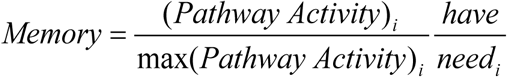

Where subscript *i* here denotes a particular learning input event. Thus, if there were not enough “Effectors Available” at the time of an event’s stimulus, *have/need* is reduced, and thus Memory is lowered.

All simulations were performed in MATLAB (The Mathworks, Natick, MA) and the code is available upon request.

---

**Supplementary Figure 1:**
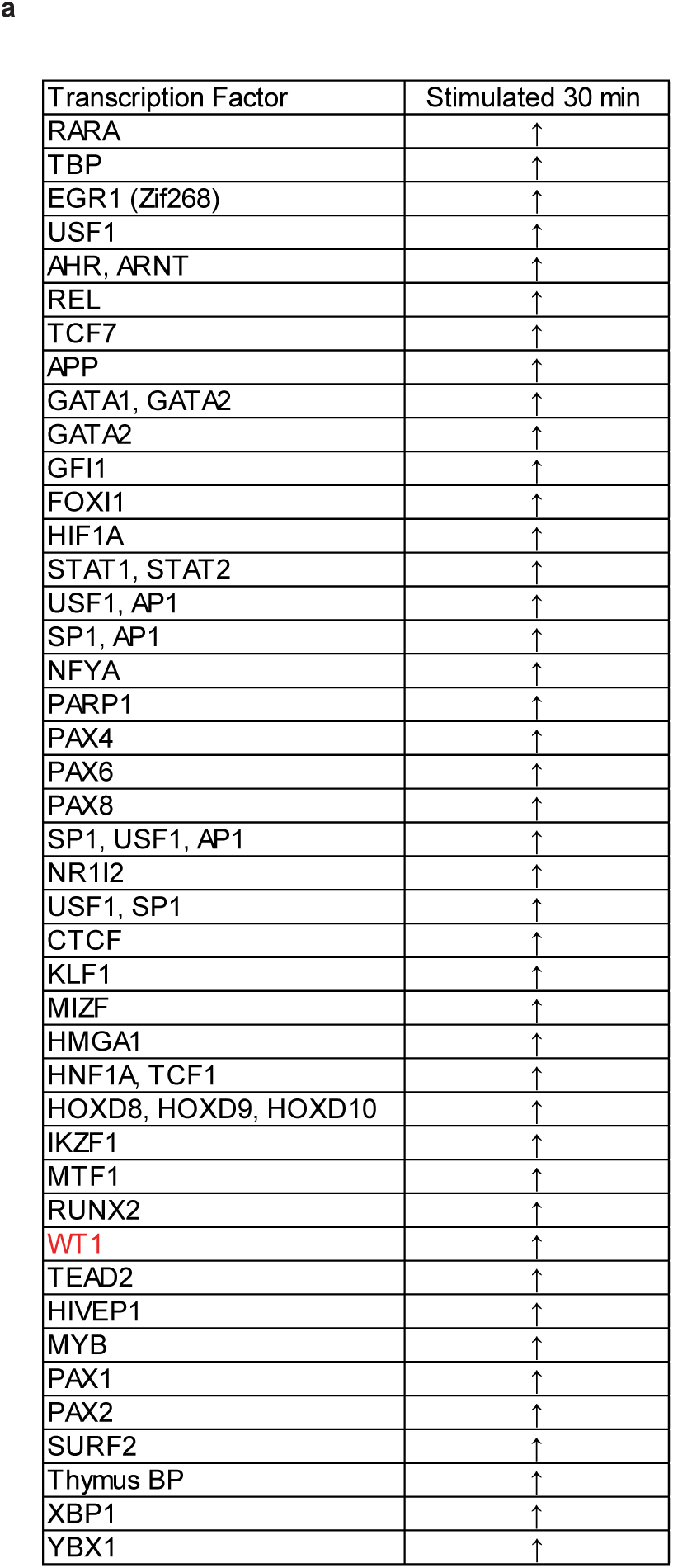
Identification of WT1 as regulator of synaptic plasticity. **a**, Summary table of transcription factors (TFs) activated 30 minutes after LTP induction. Each represented transcription factor on the table was activated in at least two separate biological replicates and with a fold change ≥ 1.3.

**Supplementary Figure 2:**
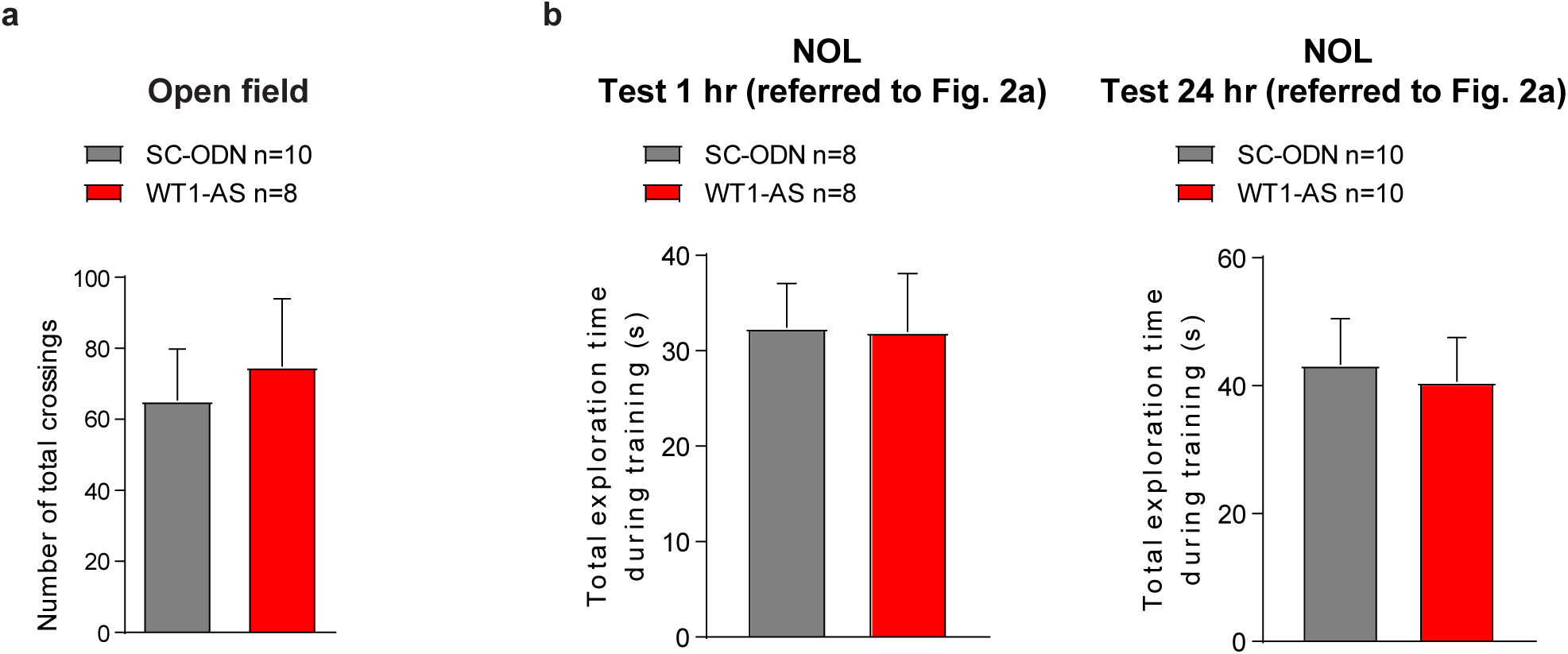
WT1 knock-down rats show normal locomotor activity and exploration during training. **a**, Injections of WT1-AS did not affect locomotion in an open field arena (data are expressed as mean ± s.e.m; unpaired t test, p>0.05). **b**, Both the WT1-AS and the SC-ODN injected groups showed no difference in the total time of exploration during training in a NOL task. The two different graphs refer to the two different time points (1hr and 24 hr respectively) reported in Fig. 2a (data are expressed as mean ± s.e.m; unpaired t test, p>0.05).

**Supplementary Figure 3:**
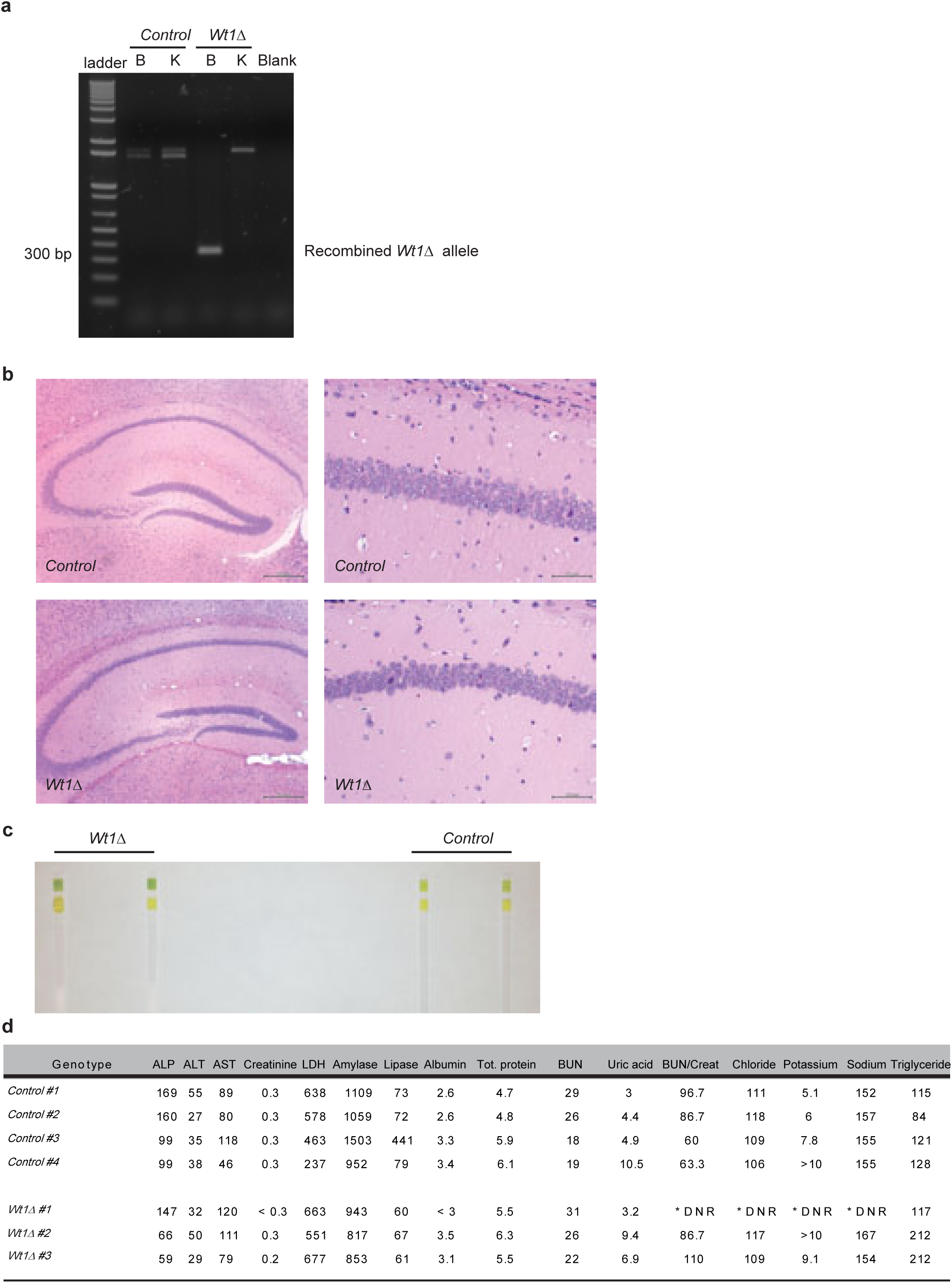
Characterization of *Wt1Δ* mice. **a,**Expression of the recombined *Wt1* allele (*Wt1 ^Δ^*) in the brain (indicated as “B”) of mice as shown by RT-PCR. Kidney (indicated as “K”) samples from both groups were negative; as expected, the recombined allele was not expressed in either tissue obtained from *Control* mice. **b**, H&E staining of adult mice hippocampal sections showed no apparent morphological differences between wild-type (*Control*) and *Wt1Δ* animals. **c**, Urine analysis indicated that kidney function was intact in *Wt1Δ* mice as shown by comparison of proteinuria levels with *Control* littermates. **d**, Blood chemistry panel of *Control* and *Wt1Δ* mice. Legend: *DNR, Did Not Report; ALP, Alkaline Phosphatase; ALT, Alanine Aminotransferase; AST, Aspartate Transaminase; LDH, Lactate Dehydrogenase; BUN, Blood Urea Nitrogen; BUN/Creat, Blood Urea Nitrogen over Creatinine Ratio.

**Supplementary Figure 4:**
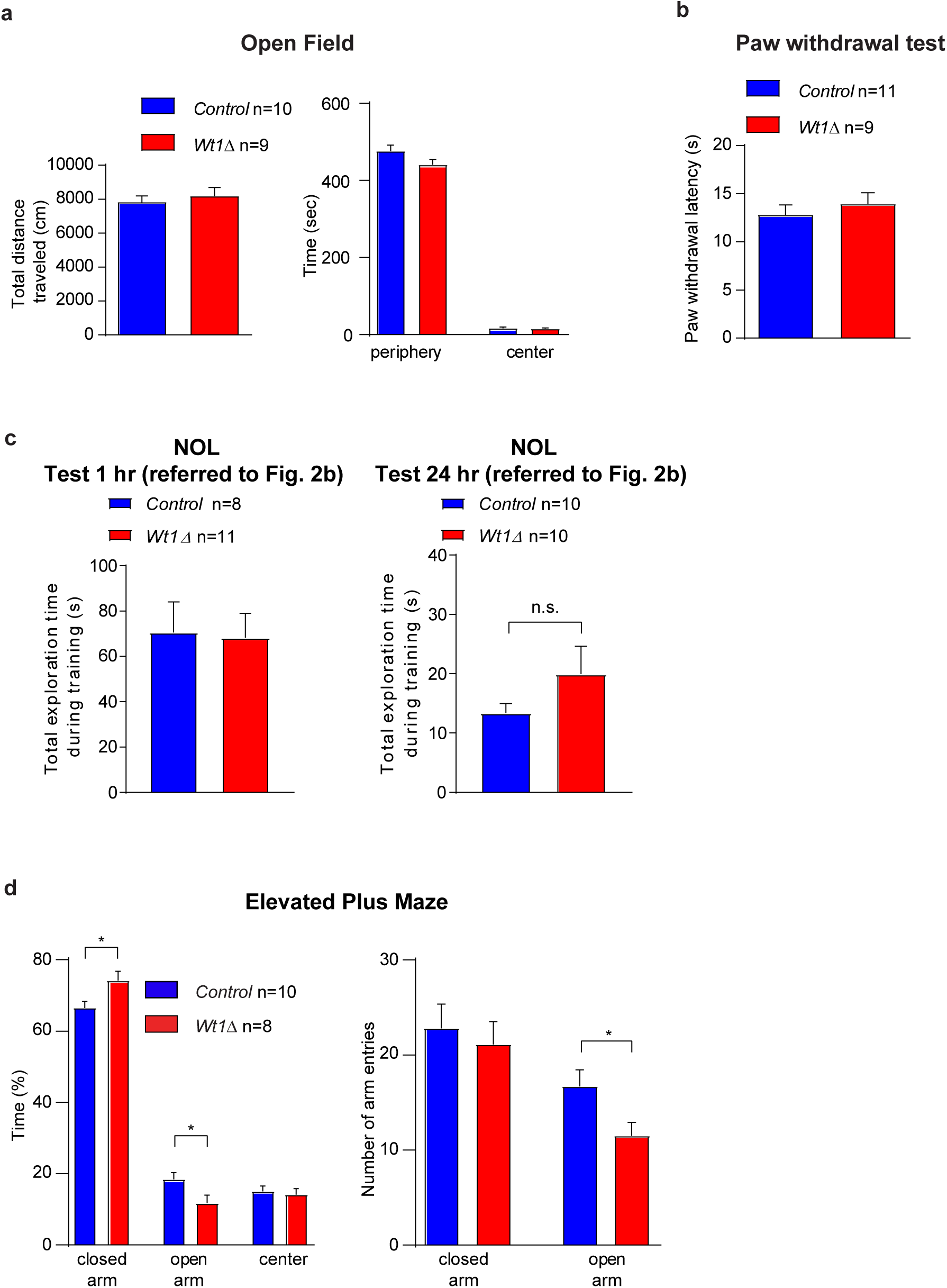
*Wt1Δ* animals show behavioral characteristics similar to their control littermates. **a**, When tested in an open field, *Wt1Δ* animals showed locomotion similar to their control littermates, expressed as total distance traveled (left panel) or time spent in the periphery or center of the open field arena (right panel: data are expressed as mean ± s.e.m.; unpaired t test, p>0.05). **b**, A paw withdrawal test was used to show that there was no difference in the nociception of *Wt1Δ* mice compared to *Control* littermates (data are expressed as mean ± s.e.m.; unpaired t test, p>0.05). **c**, *Control* and *Wt1Δ* mice showed similar total time of exploration during training in a NOL task. The two different graphs refer to the two different time points (1hr and 24 hr respectively) reported in Fig. 2b (data are expressed as mean ± s.e.m.; unpaired t test, p>0.05). **d**, Left panel: *Wt1Δ* animals spent significantly more time in the closed arm (left panel: data are expressed as mean ± s.e.m.; unpaired t test: *p=0.0249; t=2.475, df=16) and correspondently significantly less time in the open arm of the elevated plus maze compared to wild-type littermates (*Control*) (unpaired t test: *p=0.0339; t=2.320, df=16). Also, *Wt1Δ* animals entered the open arm of the maze a significant lower number of times (right panel: data are expressed as mean ± s.e.m.; unpaired t test, *p=0.0412; t=2.22, df=16).

**Supplementary Figure 5:**
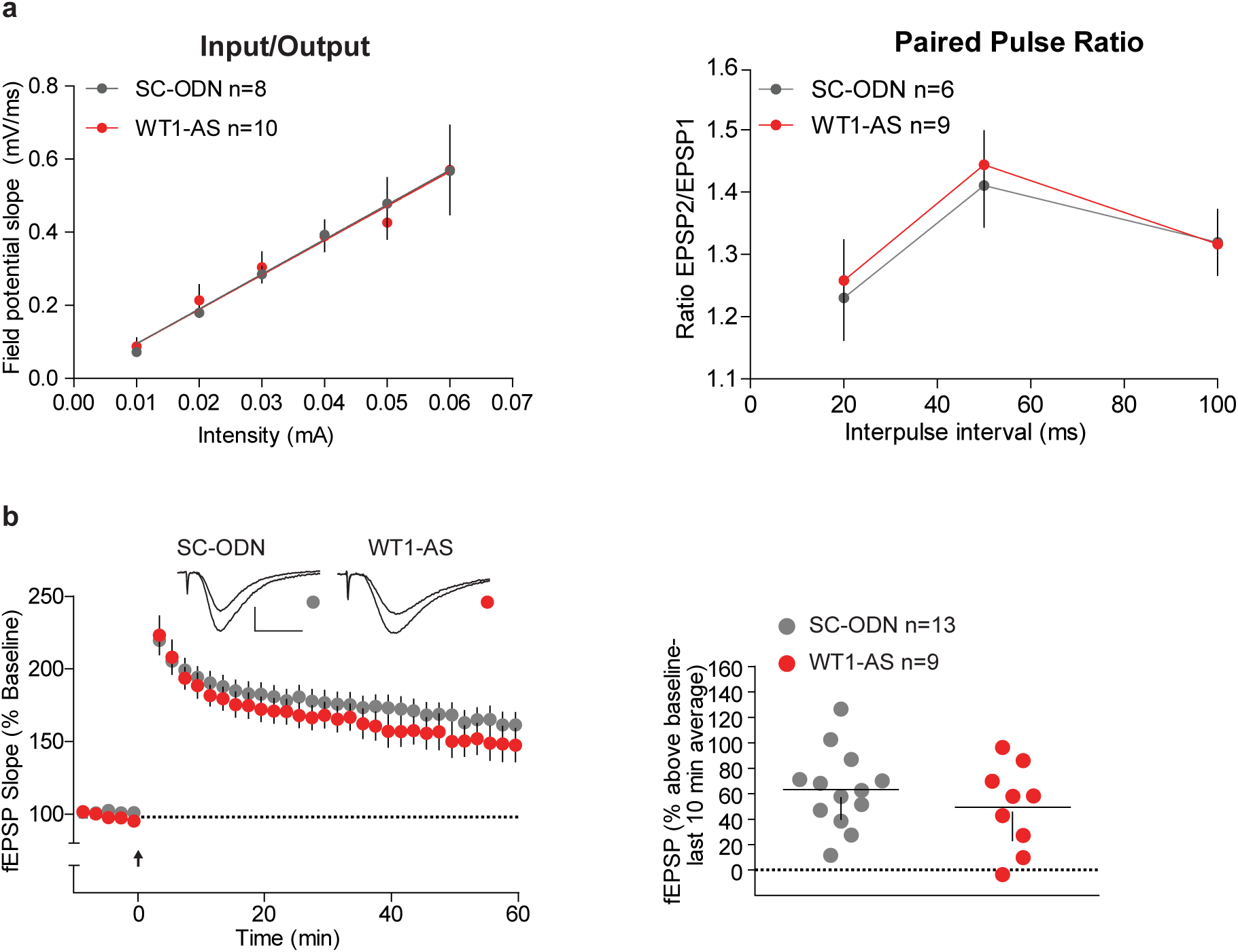
WT1 knock-down rats (WT1-AS) show basal synaptic transmission and Strong-HSF induced LTP similar to control rats(SC-ODN). **a**, Acute WT1 knock-down did not affect basal synaptic transmission measured through the analysis of the input/output relationship (left panel: data are expressed as mean ± s.e.m.; linear regression t-test, p>0.05) or the paired-pulse ratio (right panel: data are expressed as mean ± s.e.m.; two-way ANOVA RM, p>0.05) at Schaffer collateral-CA1 inputs. **b**, Injection of WT1-AS did not affect LTP induced by Strong-HFS. Representative fEPSPs show superimposed traces recorded during baseline and 60 min post-HFS. Calibrations: 0.5 mV / 10 ms. The arrow indicates time of Strong-HFS (left panel). Summary graph for the final 10 minutes of the recording (right panel: data are expressed as mean ± s.e.m.; two-way ANOVA RM: p>0.05).

**Supplementary Figure 6:**
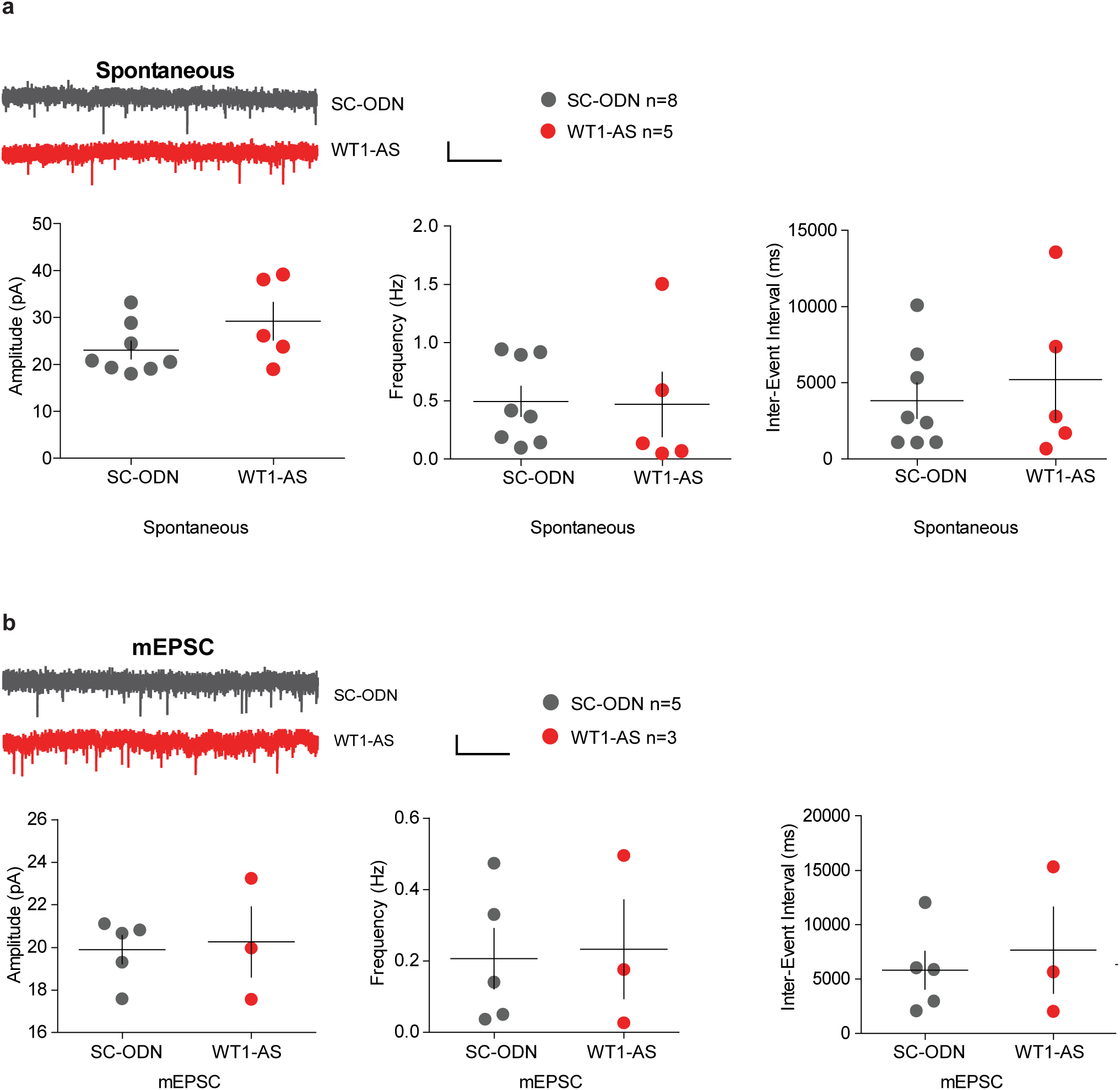
WT1 knockdown does not alter spontaneous postsynaptic currents or mEPSCs in rats. **a**, In whole-cell recordings from area CA1 pyramidal neurons of acute slices, ODN-mediated depletion of WT1 (WT1-AS) did not affect amplitude, frequency, or inter-event interval for spontaneous currents (n=8 for SC-ODN and n=5 for WT1-AS; data are expressed as mean ± s.e.m.; unpaired t tests, all p’s >1). Calibration: 20 pA / 5s. **b**, Amplitude, frequency, and inter-event interval for mEPSCs were not affected by WT1-AS treatment (n=5 for SC-ODN, and n=3 for WT1-AS; data are expressed as mean ± s.e.m.; unpaired t tests, all p’s >1). Calibration: 20 pA / 5s.

**Supplementary Figure 7:**
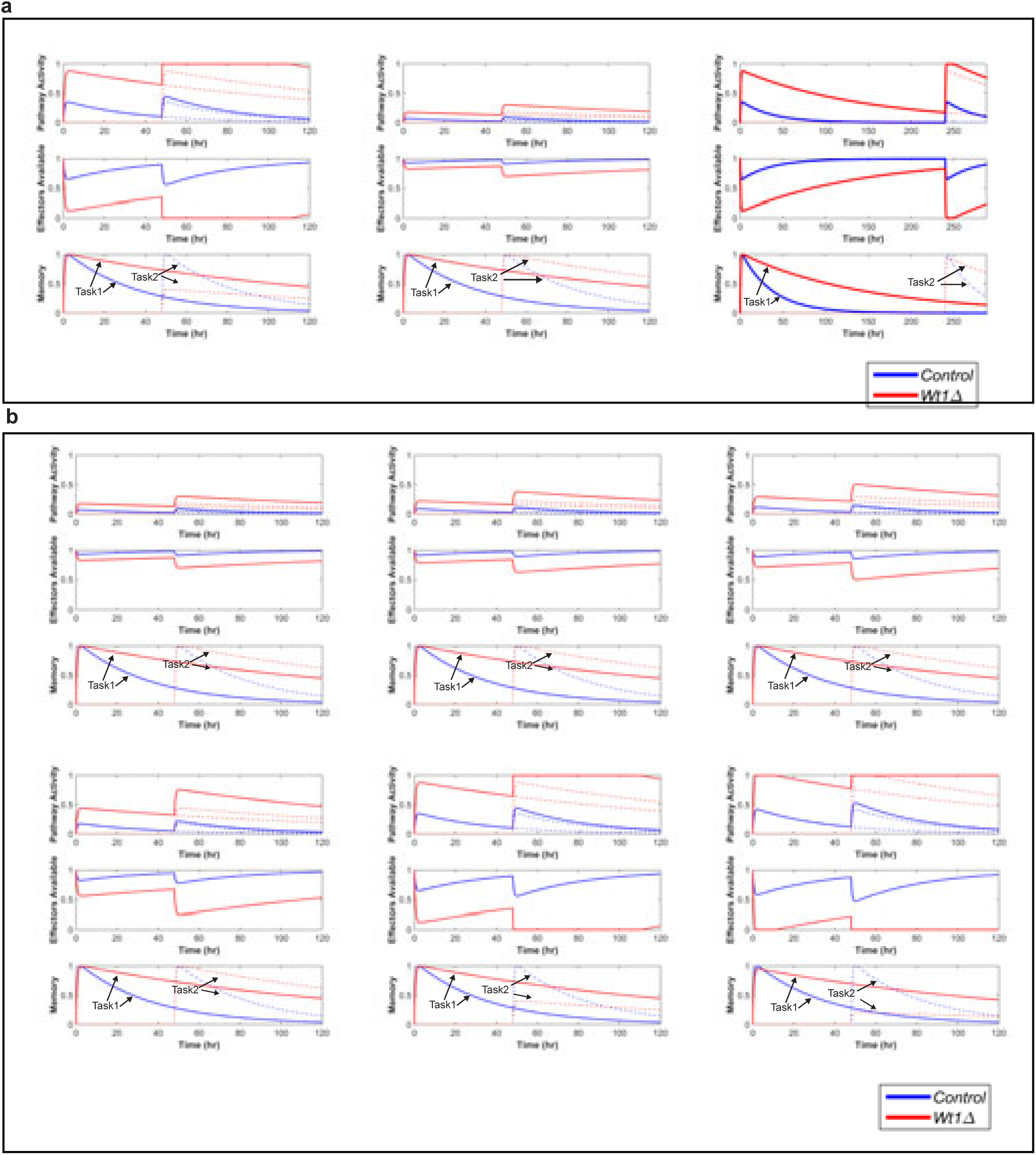
**Control theory model of WT1-mediated effect on memory flexibility. a,** Simulations using the toy control theory model. Parameter values are given in Table S1. In all panels, a first event (task1) is simulated at t=0. The strength of the input u is taken as 0.3 in the left column and the right column of panels, and as 0.05 in the middle column of panels. The second event (task2) is simulated at 48 hrs in the left and middle column of panels, and 10 days in the right column of panels. In Pathway Activity plots, the skinny dotted lines refer to contributions for individual events, and the thick solid line is the total from all events. In Memory plots, thick solid lines denote Task 1, whereas thick dashed lines denote Task 2. **b**, Effects of parameter sweeps: here the value of the input magnitude *u* was varied from 0.05 to 0.3, in increments of 0.05. As *u* is increased, the effect of saturation upon a 2^nd^ stimulation becomes evident.

**Supplementary Figure 8:**
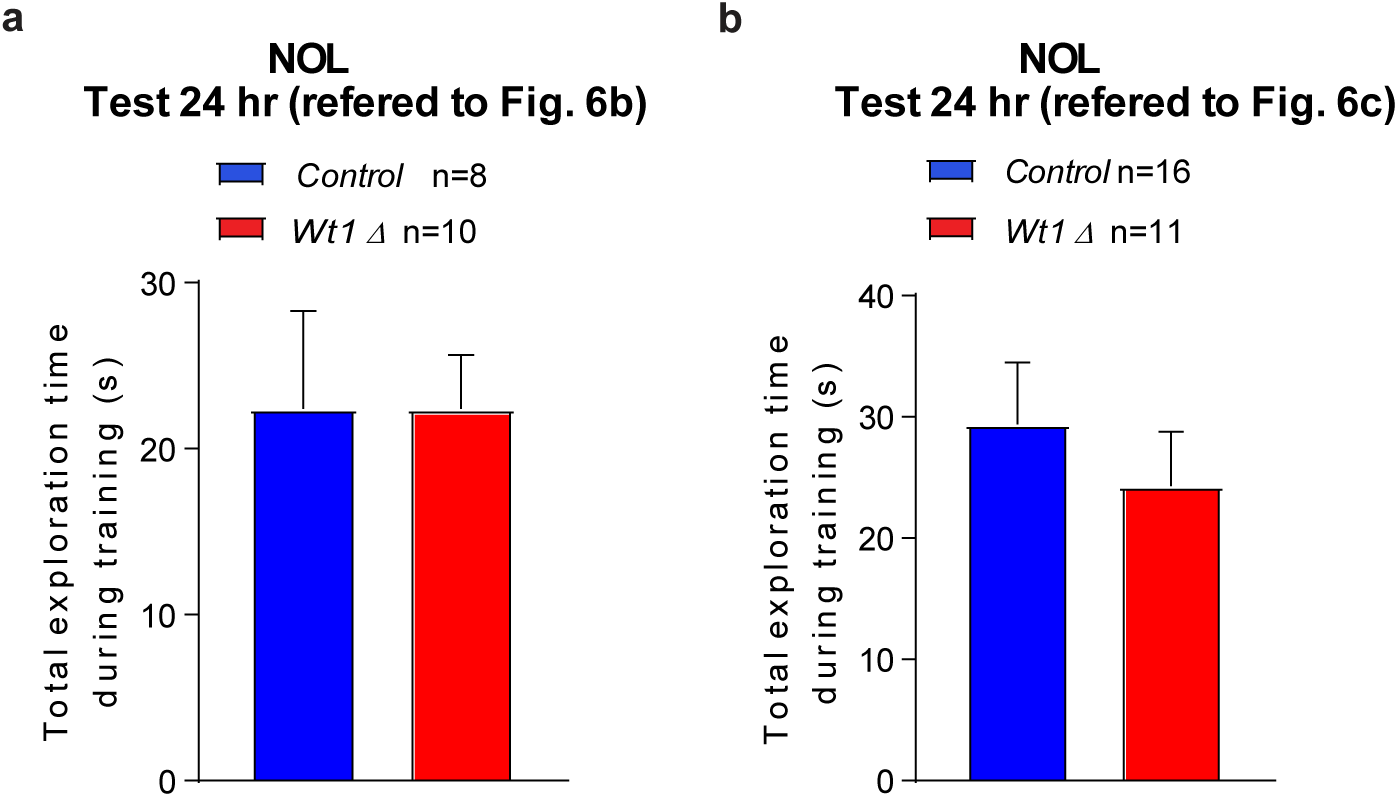
*Control* and *Wt1Δ* animals do not exhibit preference for object location during training in NOL in sequential learning protocol. **a**, Both *Control* and *Wt1Δ* animals showed no difference in the total time of exploration during training in a NOL task during a sequential learning protocol (referred to Fig. 6b; data are expressed as mean ± s.e.m.; unpaired t test, p>0.05). **b**, Both *Control* and *Wt1Δ* animals showed no difference in the total time of exploration during training in a NOL task during a sequential learning protocol (referred to Fig. 6c; data are expressed as mean ± s.e.m.; unpaired t test, p>0.05)

**Table S1:**
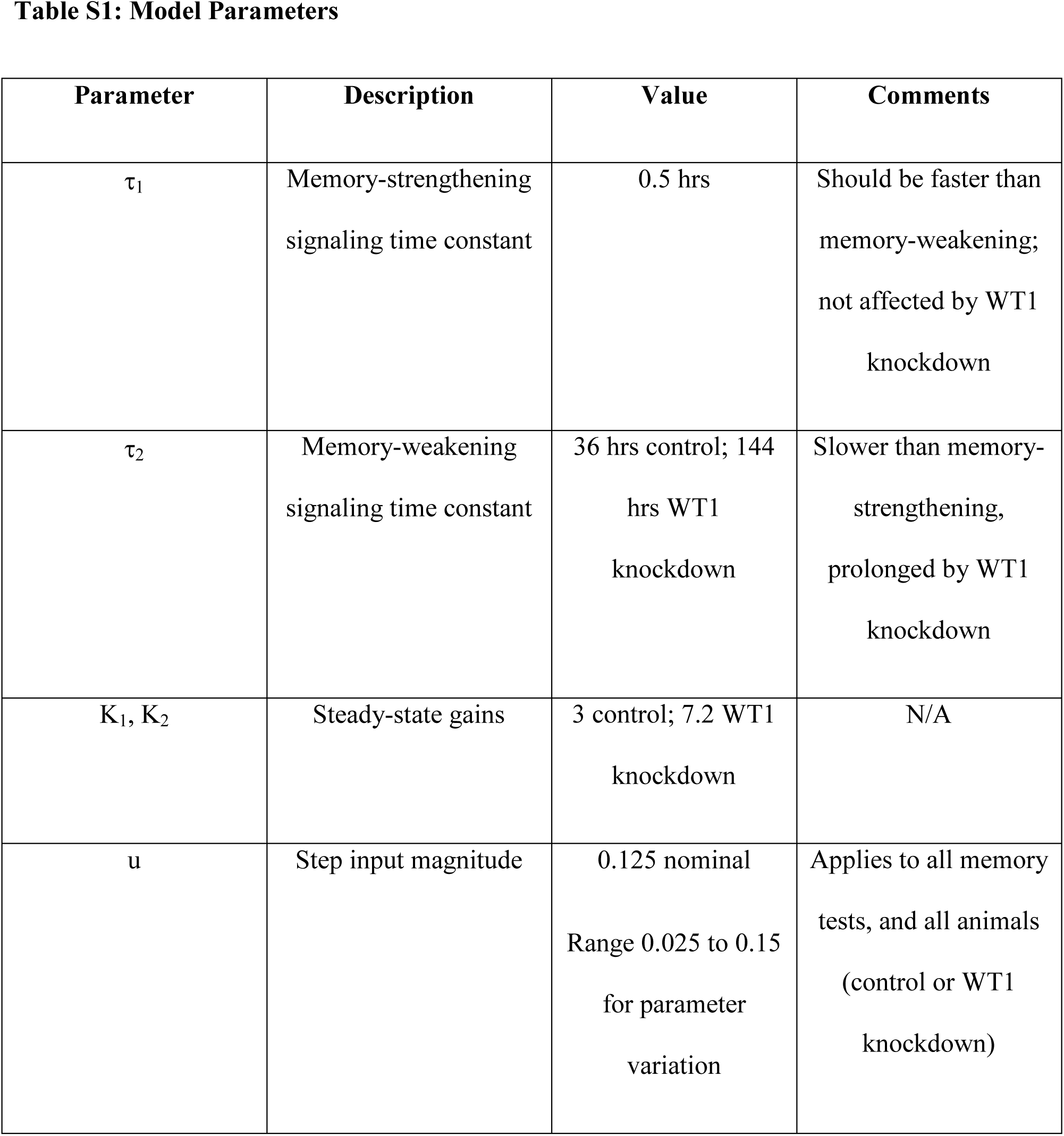
Model Parameters.

